# Connectomics Reveals a Feed-Forward Swallowing Circuit Driving Protein Appetite

**DOI:** 10.1101/2025.08.25.671815

**Authors:** Ibrahim Tastekin, Inês de Haan Vicente, Rory J. Beresford, Nils Otto, Georgia Dempsey, Scott Waddell, Carlos Ribeiro

## Abstract

Nutrient state shapes not only what animals eat, but how they eat it. We lack circuit-level mechanisms explaining how gustatory and internal state information regulates feeding at the motor control level. In *Drosophila*, protein deprivation prolongs protein-specific feeding bursts, yet the detailed motor mechanisms underlying this change remain unknown. Using EM connectomics, we identified a feed-forward pathway from protein-sensitive gustatory receptor neurons to swallowing motor neurons. At its core is the Sustain neuron, which coordinates multiple swallowing motor neurons to move food efficiently through the cibarium and pharynx. This nutrient-dependent facilitation of swallowing sustains long feeding bursts, directly linking internal state to the temporal structure of the feeding motor program. Our findings reveal how a dedicated sensorimotor circuit translates physiological need into precise motor control to drive nutrient-specific feeding appetite and highlights the power of combining EM connectomics with experimental circuit dissection.

## Introduction

Animals exhibit a rich repertoire of behaviors crucial for survival and reproduction, with feeding standing out as one of the most ancient and essential ones. From the oldest known Ediacaran organism to present-day humans, animals must maintain a balanced intake of nutrients to optimize lifespan, healthspan, and fecundity ^1–4^. Underlying this process is a sophisticated interplay of neural circuits, sensory processing, and motor control, which coordinate the acquisition, evaluation, and ingestion of food ^5–8^. Disruptions in this delicate balance, whether through nutrient deficiencies or excesses, can lead to malnutrition or metabolic diseases. In humans, an imbalanced or inadequate nutrient intake is the major cause of pathologies like scurvy and protein-energy malnutrition (PEM) ^9^, whereas their overconsumption is linked to conditions like diabetes and obesity ^10–12^. Therefore, to meet shifting physiological demands and food availability conditions, animals have evolved flexible foraging and feeding strategies ^13–15^, finely tuned by motor programs and neural networks dedicated to regulating food intake according to environmental conditions and internal states ^16–22^. Animal feeding behavior, from invertebrates to mice and humans, is implemented through a hierarchical organization of behavioral modules, collectively referred to as the feeding microstructure ^23–28^. At its lowest level, rhythmic movements of the feeding organs ^16,19,27,29^ (e.g. Drosophilae sipping with their probosces, mice licking with their tongues, or humans chewing with their jaws) mediate food intake in discrete feeding bursts. Guided by internal states, animals precisely regulate the duration and frequency of these feeding bursts to adjust overall food consumption during meals. Despite the evolutionary conservation of the higher-level feeding microstructure, the behavioral and neural processes that control the diverse low-level rhythmic motor actions to form a cohesive feeding behavior, and thereby enable nutrient balance essential for overall fitness, remain largely unknown. Analysis of feeding microstructure using edograms shows that human feeding patterns are heavily influenced by internal states (for example, deprivation or hunger level) and food palatability, leading to larger, faster meals when deprivation is prolonged ^27,30^. Similarly, in rodents, both the length and frequency of licking bursts are modulated by starvation or changes in nutritional need ^31^. In vertebrates, recent research has identified multiple nodes in the brain that adjust feeding at shortand long-timescales: from hypothalamic circuits (including AgRP and BDNF neurons) that regulate appetite and body weight via interoceptive signals ^32^, to premotor brainstem areas that integrate oral and visceral feedback to coordinate orofacial movements and ingestion ^22,33–35^. Local brainstem loops provide rapid, taste-driven feedback that curbs consumption by changing the length of licking bursts ^35^, whereas gut-derived signals relayed by vagal and hypothalamic pathways produce longer-lasting suppression of appetite ^22,32,33^. However, how these distinct brainstem centers specifically connect with gustatory circuits and the broader sensorimotor loop to collectively shape meal size and duration remains unknown. Using a high-resolution, capacitance-based feeding monitoring system (flyPAD), we previously showed that *Drosophila melanogaster* exhibits a stereotypical feeding microstructure like other animals ^25^. At its simplest, flies extend and retract their proboscis (Figure 1A) to ingest small amounts of solid food (“sipping”), which occurs in rapid (∼5 Hz), highly rhythmic bursts (Supplemental Video 1 and Figure 1B). The modulation of the frequency and duration of these feeding bursts allows flies to adapt their feeding based on physiological demands. For instance, when deprived of dietary amino acids, which are critical for lifespan and reproduction ^1^, flies specifically increase the consumption of protein-rich yeast through modular changes in the feeding structure (Figure S1): the bursts become longer and more frequent, yet individual sips remain unchanged. More specifically, amino acid-deprived females reduce the interval between bursts after 24 hours of yeast deprivation, whereas burst lengths increase when deprivation exceeds 48 hours. These findings suggest that burst frequency and duration are regulated by distinct sensorimotor and neuromodulatory pathways, although like in mammals the precise mechanisms controlling the length and frequency of feeding bursts remain largely unknown. On the sensory side of the sensorimotor circuit controlling fly feeding lies a distributed network of gustatory receptor neurons (GRNs) housed in morphologically diverse gustatory structures (Figure 1C) ^36^. We and others previously showed that Ionotropic Receptor 76b (IR76b)–expressing GRNs respond to yeast and are required for yeast appetite, with their activity being modulated by the animal’s amino acid or protein state ^8,37–39^. Consequently, IR76b-expressing GRNs function as a key entry point into the circuit governing feeding microstructure parameters underlying yeast intake. On the motor side, proboscis muscles act as the actuators for proboscis positioning and swallowing. Each is innervated by one or a few anatomically characterized motor neurons, and their associated motor actions have been extensively investigated ^17,19,36,40–42^. The intimate knowledge of the sensory and motor components of this circuit opens the opportunity to precisely dissect the higherorder circuits by which a complex, essential behavior is instantiated, coordinated, and modulated. Recent advances in high-throughput electron microscopy (EM) and automated image analysis have revolutionized our understanding of the *Drosophila* nervous system by providing near-complete connectomes at synaptic resolution ^43–51^. These advances, coupled with powerful genetic tools that enable single-cell manipulations ^52–56^, have begun to reveal how distinct sensory neuron populations and higher-order circuits cooperate to drive essential behaviors like feeding ^57,58^. In larvae, the simpler reflex of pharyngeal pumping was thoroughly mapped, illustrating how nutrient value and mechanosensory feedback can strengthen or weaken motor neuron activity ^59–61^. However, adult flies exhibit a more complex feeding repertoire involving multiple coordinated motor actions reminiscent of mammalian orofacial control. The new full adult female and male brain connectomes (FAFB – FlyWire and male CNS) ^45,62^ now offer a framework for dissecting these higher-order networks. While previous studies have identified specific gustatory receptor neurons ^38,63–72^ and key interneurons ^8,41,57,73–76^ underlying the proboscis extension response (PER) in immobilized flies, a comprehensive circuit-level picture of how these elements dynamically interact to control feeding in freely behaving animals remains elusive. Here, we combined EM connectomics, high-resolution analysis of feeding microstructure, and neurogenetic manipulations to map neurons downstream of a set of gustatory receptor neurons (dorsal taste peg GRNs) that are important for sustaining feeding bursts on yeast (a proteinaceous food of *Drosophila*) in adult flies. Connectomics analysis in this and an accompanying paper ^36^ revealed two distinct taste peg GRN types that output onto different second-order hubs that are strongly connected to motor neurons controlling proboscis positioning or ingestion. Silencing the neuronal cell types in the proboscis positioning hub did not change the length of yeast feeding bursts. Careful dissection of the ingestion hub circuits revealed the identity of a pair of interneurons (Sustain neurons) that specifically regulate the duration of yeast feeding bursts in a protein state-dependent way. We showed that Sustain neurons are strongly connected to the feeding motor neurons that are involved in swallowing and ingestion (e.g., MN11D) ^17,36,41^, but not proboscis extension. Silencing MN11D resulted in shorter feeding bursts in sucrose feeding as well, suggesting that this finding is generalizable to other food substrates. Finally, we showed that Sustain activity is necessary for controlling the ingested volume of proteinaceous food. The emerging model suggests that by anticipatorily increasing the efficiency of ingestion in each sip cycle via the Sustain neuron, dorsal taste peg GRNs allow the fly to increase the duration of feeding bursts and therefore yeast intake. Combined with the recent findings in mice, our work highlights the role of nutrient state-dependent swallowing efficiency as a key motor action in the control of the feeding microstructure.

**Figure 1.**
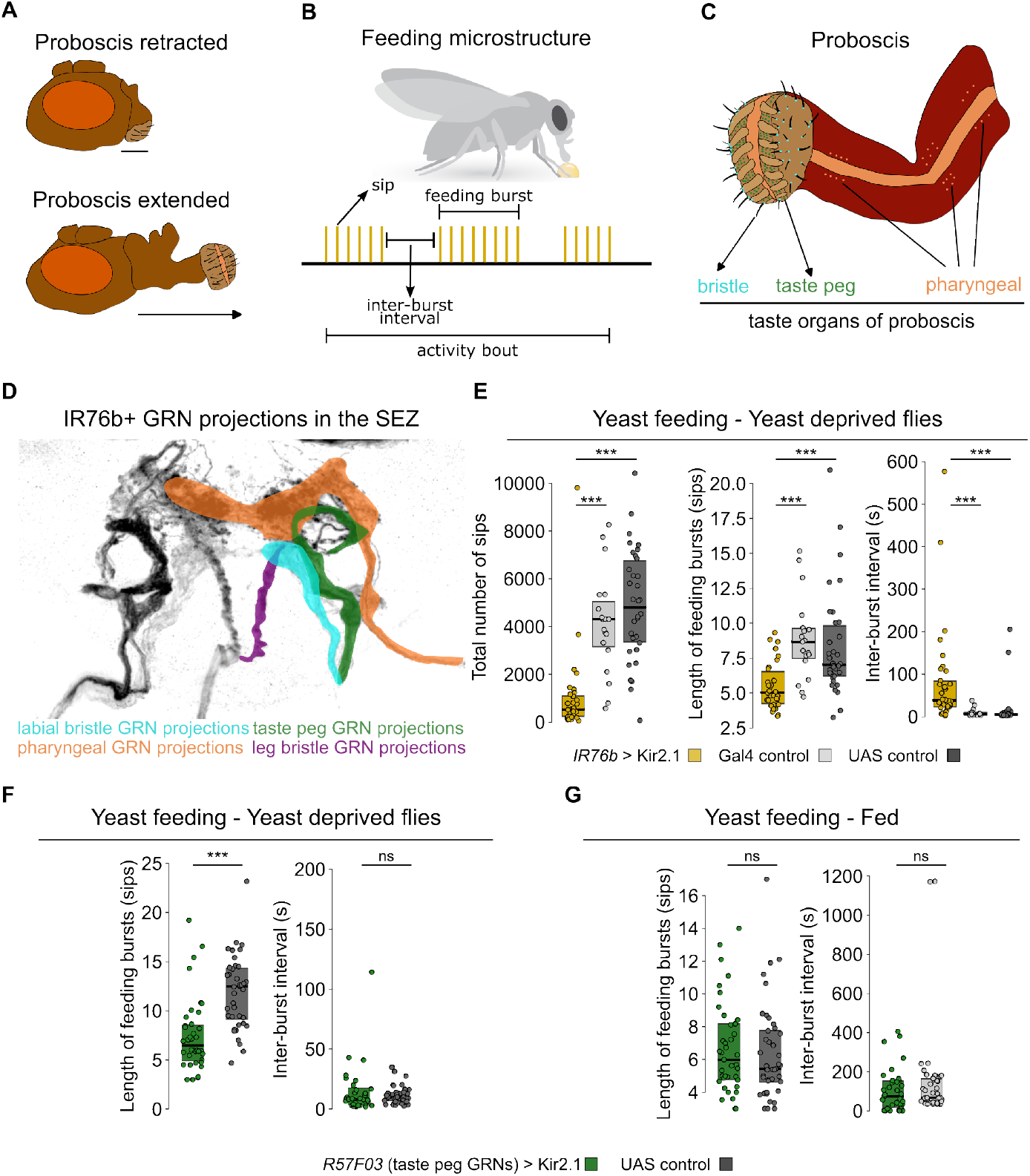
Modulation of feeding microstructure parameters by distinct IR76b positive gustatory receptor neuron populations. **A.** Schematic representation of the proboscis positions during feeding. Retracted (top panel) and extended (bottom panel) positions are depicted. **B**. The feeding microstructure of *Drosophila melanogaster*. An extension-ingestion-retraction cycle of the proboscis is called a ‘sip’. Sips are organized into feeding bursts with stereotypical sip durations and inter-sip intervals. Multiple feeding bursts form an activity bout ^25^. **C**. Taste organs of the proboscis. Hair-like bristle sensilla (cyan) are found on the outer surface of the labellum, and taste peg sensilla (green) reside in the inner surface of the labellum. Three pharyngeal sense organs (light orange) are found along the pharynx, located within the proboscis. **D**. The projections of IR76b expressing gustatory receptor neurons (GRNs) in the subesophageal zone (SEZ) of the fly brain. Different sensillar projections are color-coded, as in C. **E**. The effect of silencing IR76b-expressing neurons on yeast feeding. To silence IR76b neurons, the inward rectifying potassium channel Kir2.1 was expressed under the control of *IR76b-Gal4* in yeast-deprived flies. Silencing IR76b neurons led to a drastic decrease in the total number of sips (left), the length of feeding bursts (middle), and an increase in the duration of inter-burst intervals (right) when compared to the UAS control: *w1118* x *UAS-Kir2*.*1, tubGal80ts*, and the Gal4 control: *IR76b-Gal4* > *w1118*. **F**. and **G**. The effect of silencing taste peg GRNs using *R57F03-Gal4*, which labels taste peg neurons, on yeast feeding. Yeast-deprived animals in F, and fully-fed flies in G. Inter-burst intervals were not affected (right panels). The length of feeding bursts is affected upon silencing **F**. in yeast-deprived but **G**., not in fully fed flies (left panels). **E-G**. The conditional expression of Kir2.1 was controlled using a ubiquitously expressed temperature-sensitive Gal80 (*tubulin-Gal80ts*). The flies were moved to 30^*°*^ C one day before the experiment to switch on the expression of Kir2.1. Filled circles indicate individual flies. Boxes indicate the interquartile ranges. Horizontal lines indicate the median value. ns: not statistically significant, Wilcoxon rank-sum test. P-values are indicated. ***: *p <* 0.001

## Results

### Modulation of feeding microstructure parameters by distinct IR76b positive gustatory receptor neuron populations

IR76b is expressed in GRNs located in different types of taste sensilla: bristle sensilla on the external surface of the labial palps, taste peg sensilla on the interior part of the labial palps, pharyngeal sensilla (Figure 1C), and bristle sensilla on the legs ^37,39,77^. They all project their axons to the gustatory neuropil of the fly brain, namely the subesophageal zone (SEZ) (Figure 1D). We have previously shown that mating and amino acid deprivation synergistically lead to an increase in yeast feeding ^8,13,39^. Silencing all IR76b neurons using an *IR76b-Gal4* transgene ^39^ in mated and yeast-deprived female flies led to a severe reduction in the total number of yeast sips (Figure 1E, left panel). This phenotype is characterized by an inability to increase the length of feeding bursts on yeast (Figure 1E, middle panel) and to reduce the duration of inter-burst intervals (Figure 1E, right panel). However, as we showed before ^39^, silencing only taste peg GRNs (tpGRNs) using *R57F03-Gal4* driver line led to a decrease in the length of yeast feeding bursts in mated flies deprived of yeast for five days (Figure 1F, left panel). Interestingly, tpGRNs were dispensable for modulating the length of the yeast feeding bursts in fully fed flies (Figure 1G, left panel). Moreover, silencing tpGRNs did not affect the inter-burst intervals in either yeast-deprived or fully fed flies (Figure 1F and G, right panels). These findings suggest that the sensorimotor pathway downstream of tpGRNs is important for modulating specifically the length of feeding bursts in response to the changes in the fly’s nutrient state. Therefore, taste peg GRNs serve as an ideal entry point to study sensorimotor circuit mechanisms underlying nutrient-state-dependent and microstructure parameter-specific (i.e., length of feeding bursts) motor programs.

### Taste peg gustatory receptor neurons in a female connectome

To map the sensorimotor connectome downstream of tpGRNs at synapse resolution, we decided to make use of a publicly available electron microscopy reconstruction dataset (FAFB – FlyWire) ^44^. We identified and proofread the SEZ projections of the entire tpGRN population through the labial nerves in this dataset, revealing 37 tpGRNs on each hemisphere (Figure 2A and B). These numbers are consistent with what had been described using light and transmission electron microscopy (42 ± 4 tpGRNs on each labial palp) ^78^. Next, we performed morphological clustering using NBLAST comparisons ^79^ of tpGRNs to test whether there are different tpGRN morphotypes. Our analysis revealed two main tpGRN morphotypes in both hemispheres (Figure 2CF). Most tpGRNs comprised a group that we named “claw tpGRNs” (ctpGRNs) (Figure 2C, D and F, magenta) due to lateral claw-like projections (Figure 2F, arrows). A much smaller number of tpGRNs formed another cluster that we named “dorsal tpGRNs” (dtpGRNs) (Figure 2C-E, blue) due to their characteristic dorsal projections (Figure 2E, arrows). Because of technical reasons, we could not complete the proofreading of one and two incomplete tpGRNs on the left and right hemisphere, respectively (Figure 2C and D, Figure S2). These incomplete tpGRNs cluster with ctpGRNs in terms of connectivity (see Figure 3). Therefore, we included them in the ctpGRN cluster. However, these incomplete neurons might also be taste peg mechanosensory neurons (tpMSNs) as both tpGRNs and tpMSNs follow the same tract ^80^. Morphological clustering, therefore, suggests that there are two distinct types of tpGRNs in *Drosophila*.

**Figure 2.**
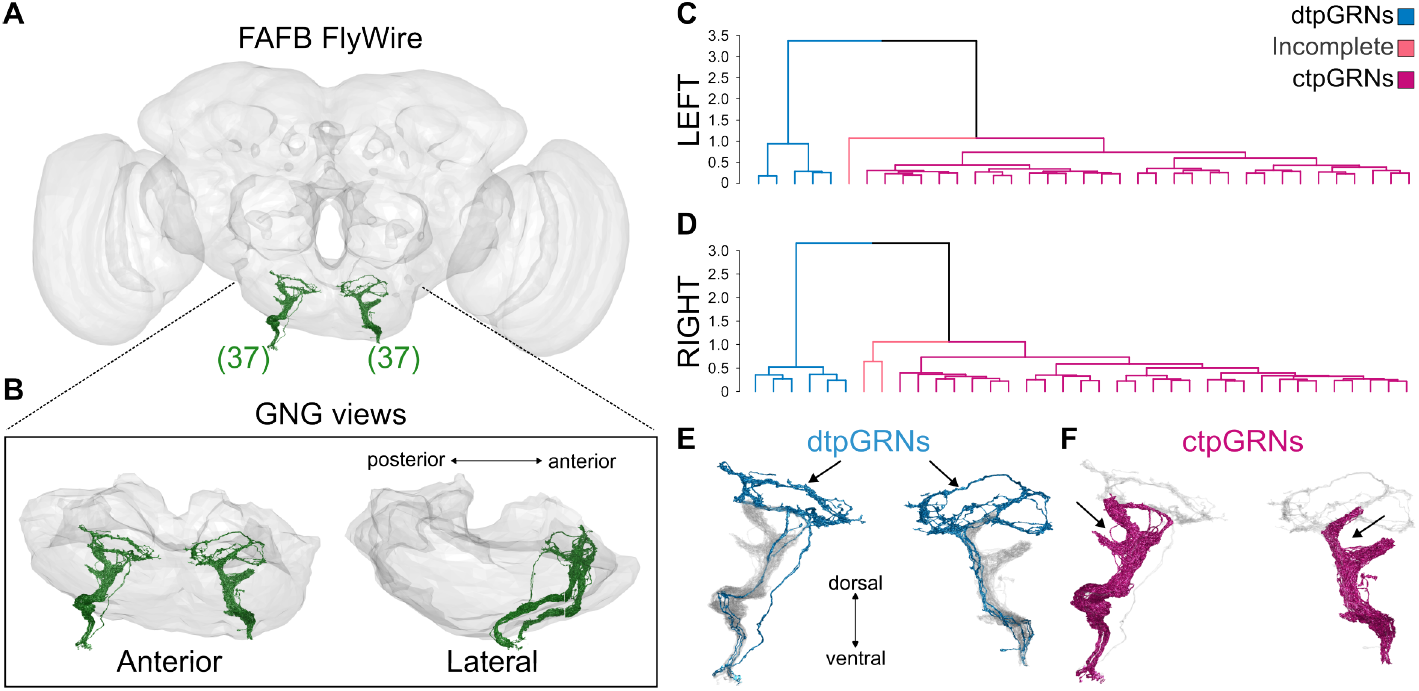
Connectomics reveals that taste peg GRNs comprise two morphologically distinct subtypes. **A.** The taste peg gustatory receptor neuron (tpGRN) projections (green) in the electron microscopy volume, FAFB – FlyWire. The brain neuropile is shown as a semi-transparent mesh (anterior view). Numbers in brackets indicate the number of identified tpGRNs on each side of the brain. **B**. Close-up view of tpGRN projections in the gnathal ganglia (GNG) of the SEZ. Anterior (left) and lateral (right) views are shown. The GNG is shown as a semi-transparent mesh. **C** and **D**. Hierarchical morphological clustering of taste peg GRNs based on computed NBLAST scores. **C**. Left and **D**. right hemisphere. In both hemispheres, two main clusters were identified: dtpGRNs: dorsal tpGRNs (blue), ctpGRNs: claw tpGRNs (magenta). A third cluster was termed incomplete: it contains tpGRNs of which proofreading could not be completed due to technical reasons (pink). **E**. Morphology of dtpGRNs (blue, anterior view). Arrows indicate the characteristic dorsal loop of these neurons. ctpGRNs are plotted in light grey. **F**. Morphology of ctpGRNs (magenta, anterior view). Arrows indicate the claw-like lateral projections of these neurons. dtpGRNs are plotted in light grey.

**Figure 3.**
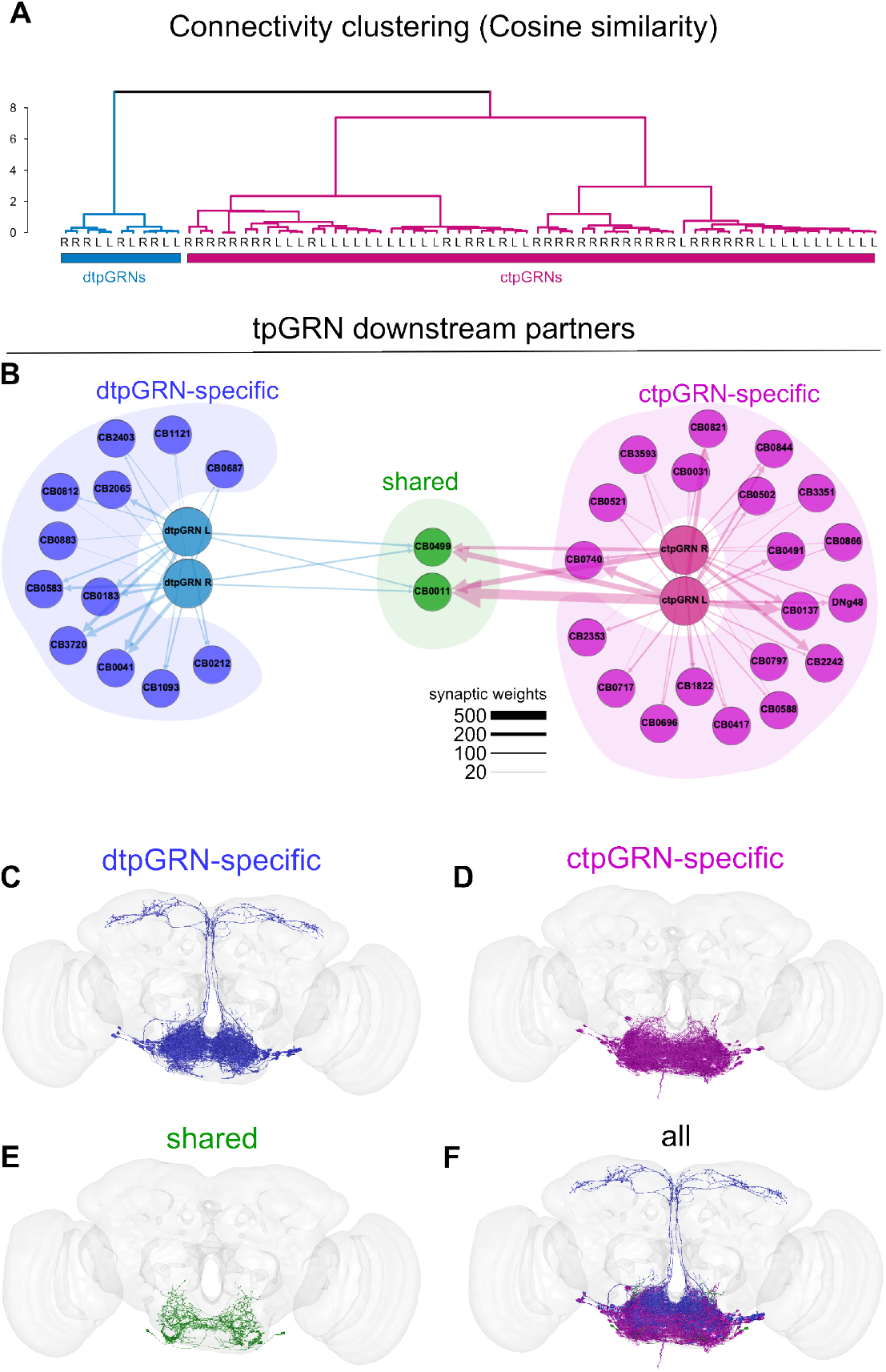
Cell typing the taste peg GRNs. **A.** Hierarchical clustering of tpGRNs using pairwise cosine similarity scores based on downstream connectivity. dtpGRNs (blue) and ctpGRNs (magenta) form two distinct clusters independently of the side to which they project. L: tpGRN with a left-hemisphere projection, R: tpGRN with a righthemisphere projection. **B**. tpGRN downstream connectivity. The Yifan Hu’s proportional network layout method is used to plot the downstream network from dtpGRNs (light blue circles) and ctpGRNs (light magenta). dtpGRN-specific (dark blue), ctpGRN-specific (light purple), and shared (green) downstream neuronal cell types with more than 19 synapses with either tpGRN type on each hemisphere are shown. The thickness of the arrows is proportional to the synaptic weight between cell types. The FlyWire cell type of each node is indicated. **C-F**. Morphological representation of downstream neurons in FAFB – FlyWire. **C**. dtpGRN-specific downstream partners, **D**. ctpGRN-specific downstream partners, **E**. shared downstream partners, and **F**. overlay of all types. Anterior view. The brain neuropile is shown as a semi-transparent mesh.

### Downstream connectivity of tpGRNs reveals functional segregation of tpGRN subtypes

If ctpGRNs and dtpGRNs are indeed two functionally distinct populations of GRNs, they would be expected to diverge in their synaptic connectivity patterns. To test this, we computed a cosine similarity matrix between pairs of tpGRNs using synaptic weights between each tpGRN and all tpGRN postsynaptic partners that have at least 2 synapses with any of the tpGRNs (Figure S3). Hierarchical clustering of cosine similarity as well as Euclidean distances based on absolute synaptic weights with downstream partners separated dtpGRNs and ctpGRN morphotypes into two different connectivity clusters (Figure 3A, Figure S3 and Figure S4A). Moreover, the highest average silhouette width was obtained with k=2 clusters (Figure S4B), reinforcing the finding that dtpGRNs and ctpGRNs are two distinct populations which have different postsynaptic connectivity patterns. In an accompanying study using the male CNS EM dataset, we have shown that male flies also have both types of tpGRNs ^36^, highlighting the reproducibility of this finding.

Further analysis of downstream connectivity with either tpGRN population (a cumulative synaptic weight threshold of > 19 for the entire dtpGRN or ctpGRN population) revealed that there are three main groups of tpGRN postsynaptic partners in terms of their tpGRN connectivity: postsynaptic neurons that receive inputs from only dtpGRNs (dtpGRN-specific), only ctpGRNs (ctpGRN-specific), and both tpGRN types (shared) (Figure 3B, Figure S5 and S6). While shared and ctpGRN-specific postsynaptic neurons are almost exclusively local SEZ neurons (Figure 3D-F, Figure S5 and S6), dtpGRN-specific population comprised both local SEZ neurons and SEZ output neurons (SEZONs) that ascend to the superior neuropils (SNP) (Figure 3C and F, Figure S5 and S6). Except for one SEZON, which is predicted to express serotonin, each downstream cell type was predicted to express one of the fast-acting neurotransmitter types: acetylcholine, glutamate, or GABA (Figure S6). As both GRN projections and dendrites of feeding motor neurons are confined within the SEZ ^19,36,50^, we propose that local SEZ neurons in all three groups might be involved in sensorimotor transformations underlying different aspects of feeding motor control. SNP projecting SEZONs downstream of dtpGRNs might contribute to higher-order computations because the SNP region is densely innervated by neuronal cell types controlling navigation ^81^, foraging decisions ^15^, neurosecretion ^82^, learning and memory ^83–85^. The descending neuron cell type downstream of ctpGRNs might contribute to the coordination of feeding with locomotor activity ^86–88^. Here, we focused on dissecting the local SEZ circuits to unravel sensorimotor mechanisms underlying the nutrient state-specific modulation of the length of feeding bursts.

### Connectivity of tpGRN postsynaptic partners with proboscis motor neurons reveals two hubs of sensorimotor control

The proboscis consists of three parts termed the labellum, haustellum, and rostrum ^19,42^. During *Drosophila* feeding, sips are generated by the coordinated activity of muscles controlling proboscis positioning (the movements of rostrum, haustellum, and labellum) and swallowing (Figure 4A), leading to an extension – food ingestion – retraction cycle ^19,25,36,42^. The food ingestion step can be further subdivided into the following steps: (1) labial palps open (labellar spreading), (2) food is sucked in, (3) pumped through the pharynx into the cibarium (pharyngeal cavity), and (4) enters the esophagus to move towards the gut ^17,41,89^. It was shown that the activity of pharyngeal muscles is important for the rate and amount of ingestion ^17^. On solid food, the proboscis is retracted upon ingestion, followed by the subsequent proboscis extension – ingestion – proboscis retraction cycles leading to a feeding burst ^25^.

**Figure 4.**
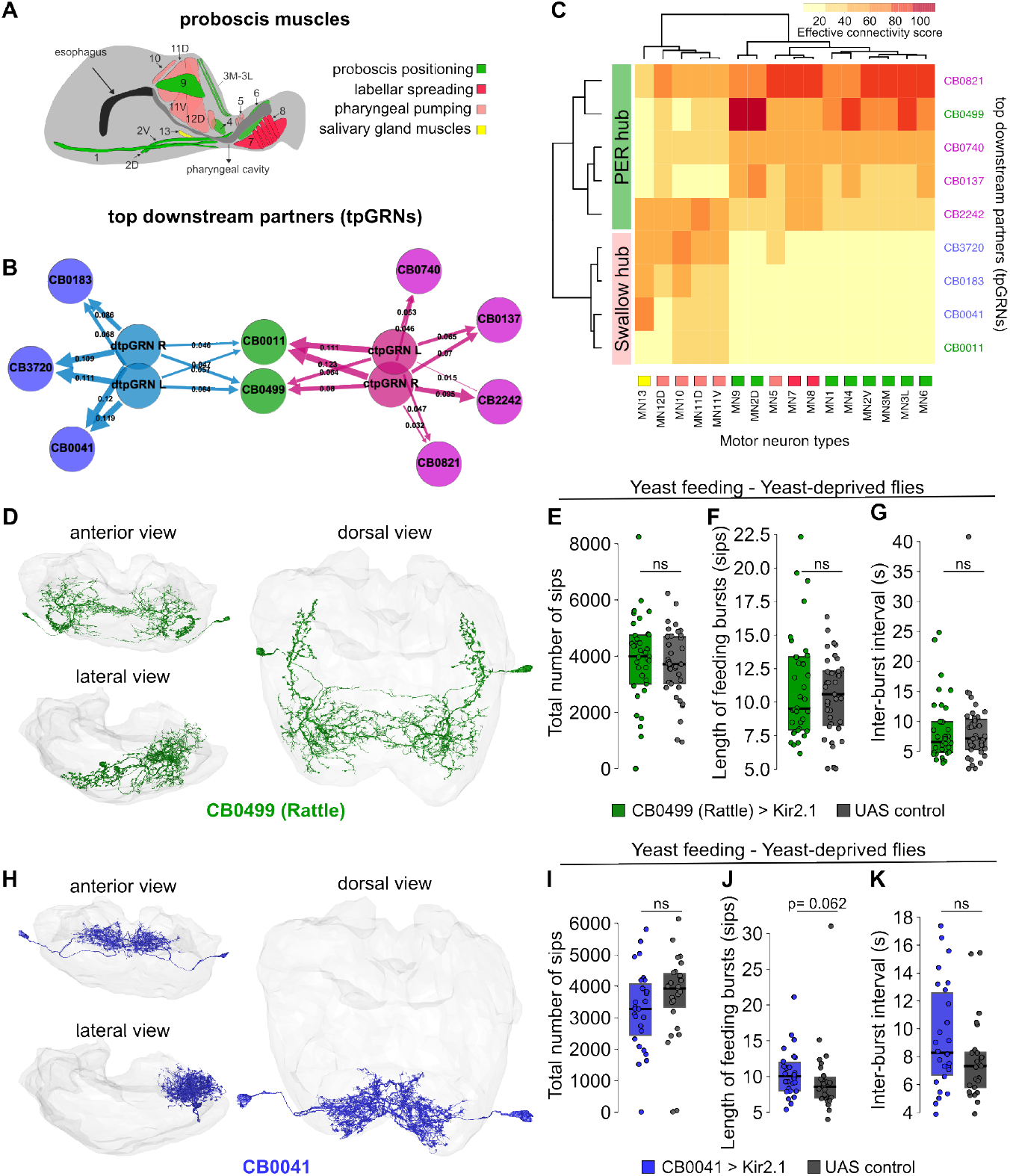
Effective connectivity analysis reveals two hubs of tpGRN downstream neurons with distinct connectivity to proboscis motor neurons. **A.** Proboscis muscle groups responsible for the movement of the proboscis during proboscis extension-retraction (labellum, haustellum, and rostrum) and ingestion (pharyngeal pumping and salivary gland). Muscle groups are color-coded according to their function. **B**. Top downstream partners of dtpGRNs and ctpGRNs. Criteria for selecting top partners are explained in the main text. Proportional synaptic strengths from tpGRNs to downstream partners are shown as numbers on the arrows. The thickness of the arrows is proportional to the synaptic weight between cell types. L: left, R: right. **C**. Heatmap and hierarchical clustering of top tpGRN downstream partners using effective connectivity scores with different motor neuron types. Motor neurons are color-coded according to their function. Right: tpGRN downstream partners are color-coded according to their tpGRN connectivity (blue: dtpGRN-specific, magenta: ctpGRN-specific, and green: shared). Hierarchical clustering reveals two downstream hubs: A PER hub (green) and a Swallow hub (salmon). **D**. Morphological representation of CB0499 (also known as Rattle) from different points of view. The GNG is shown as a semi-transparent mesh. **E-G**. Silencing CB0499, which is in the PER hub, using Kir2.1 does not influence the feeding microstructure parameters for yeast feeding in yeast-deprived flies. **H**. Morphological representation of the dtpGRN-specific downstream partner CB0041 from different points of view. The GNG is shown as a semi-transparent mesh. **I-K**. Silencing the Swallow Hub downstream neuron CB0041 in yeast-deprived flies leads to an almost statistically significant increase in the length of yeast feeding bursts (middle panel, p-value is indicated). Filled circles indicate individual flies. Boxes indicate the interquartile ranges. Horizontal lines indicate the median value. UAS control: *Empty Split-Gal4 x UASKir2*.*1*. ns: not statistically significant, Wilcoxon rank-sum test.

Which motor neurons, and thus which muscle movements, are controlled by tpGRNs to regulate the length of feeding bursts? To answer this question, we first identified the FAFB – FlyWire instances corresponding to the effectors of feeding, the motor neurons (MNs) innervating different proboscis muscles, described in detail by McKellar et al. ^19^ (except MN12V, for which we could not identify a FAFB instance) (Figure S7). MN cell types were further cross-validated using the newly released male CNS dataset ^36^. We then set out to map the connectivity of the top postsynaptic partners of the dtpGRN-specific, ctpGRN-specific, and shared groups of downstream neurons to the proboscis motor neurons. The top partners for each group were defined as the postsynaptic cell types with percentage synaptic weights exceeding the 95th quartile of the distribution of tpGRN output synapses for at least half of the neurons within the same cell type (Figure S8, note that some cell types have more than one neuron per hemisphere). This criterion led to nine postsynaptic neuronal cell types: four ctpGRN-specific, two shared, and three dtpGRN-specific cell types (Figure 4B). To identify to which motor neurons these different second-order neurons connect to, we used a computational strategy called effective connectivity (see Methods for details) ^87^. Briefly, effective connectivity computes the connectivity strength of any given pair of neurons in the connectome, including monosynaptic as well as polysynaptic connections, normalized by the total flux through these neurons. For polysynaptic connections, we restricted our analysis to eight layers since tpGRN downstream partners reach a plateau for the number of neurons they can be connected to after 6-9 hops (synapse threshold = 5, data not shown). We then used the effective connectivity scores between the tpGRN postsynaptic neuron–proboscis MN pairs to hierarchically cluster the tpGRN second-order neurons in relation to the proboscis MN population (Figure 4C). This analysis revealed two clusters of tpGRN postsynaptic partners. The first cluster consisted of four ctpGRN-specific (CB0137, CB0740, CB0821, and CB02242) and one shared (CB0499) tpGRN postsynaptic partner (Figure 4C).

This cluster was predicted to be more strongly connected to the motor neurons that control rostrum, haustellum, and labellum movements ^19,42^ compared to the other cluster (Figure 4B and C). This cluster contains the FlyWire cell type CB0499 (also known as the Rattle neuron) that has previously been shown to be necessary to promote proboscis extension reflex (PER) towards a drop of sugar solution in immobilized flies via MN9, and sufficient to trigger proboscis extension upon acute optogenetic activation ^57^. We therefore termed this cluster the “PER hub”. As expected, CB0499 has relatively high effective connectivity with MN9 and MN2D (Figure 4C), both of which are involved in rostrum movements underlying proboscis extension ^19^. This finding suggests that the effective connectivity approach was successful in predicting known functional roles of specific gustatory circuit neurons. Surprisingly, the neurons comprising the second cluster were predicted to be poorly converging onto the motor neurons that control proboscis extension and retraction, while preferentially connecting to the pharyngeal pumping motor neurons, which are involved in swallowing (Figure 4C) ^17,19^. This cluster comprised three dtpGRN-specific (CB0041, CB0183, and CB3720) and one shared (CB0011) tpGRN postsynaptic partners. Based on this effective connectivity pattern, we termed this cluster the “swallowing hub”. Interestingly, ctpGRN-specific postsynaptic neurons are found only in the PER hub, while dtpGRN-specific neurons were only found in the swallowing hub, reinforcing the hypothesis that these neurons may serve distinct functions (Figure 4C).

### The tpGRN PER hub is dispensable for modulating the length of feeding bursts

Which of these two hubs could facilitate longer feeding bursts on yeast in yeast-deprived flies? One simple and intuitive hypothesis would be that the sensorimotor circuit downstream of tpGRNs acts on the proboscis motor neurons mediating proboscis retraction-extension movements to add more sips to feeding bursts. To test this hypothesis, we decided to neurogenetically manipulate the function of CB0449 (Rattle ^57^) neurons in the PER hub and test for phenotypes in the feeding microstructure in freely behaving animals. CB0499 is a single pair of local SEZ neurons (Figure 4D) that receives input from sweet-sensing GRNs of labial bristle sensilla as well as both types of tpGRNs (Figure 4B and Figure S9). Surprisingly, silencing CB0449 in female flies by expressing Kir2.1^90^ using a specific Split-Gal4 driver line (*Rattle Split-Gal4* > *Kir 2*.*1*) had no effect on the feeding microstructure towards yeast in freely behaving, yeast-deprived flies (Figure 4E-G). This included the total amount of feeding (Figure 4E), the length of yeast feeding bursts (Figure 4F), and the inter-burst intervals (Figure 4G). In contrast, when we analyzed the phenotype for sugar feeding in yeast-deprived flies, CB0499 silencing had a small but significant effect on inter-burst intervals, while having no effect on the length of feeding bursts and total number of sips (Figure S10A-C). These findings suggest that CB0499-driven PER is dispensable for modulating the length of feeding bursts on either yeast (Figure 4F) or sucrose (Figure S10B), but important for initiating sugar feeding bursts (Figure S10C), a result which is consistent with findings obtained earlier in immobilized flies (PER assay) ^57^. To strengthen this finding, we silenced the Fdg (CB0038) ^41^, Bract (DNge173 and DNge174), and Roundup (CB0553) neurons, which this analysis as well as previous work on sugar GRNs ^57^ have shown to be downstream of CB0499 (Rattle) and upstream of the PER-triggering motor neuron MN9 (Figure S10D). Similar to what we found for CB0499, none of these neurons was necessary for modulating the length of yeast feeding bursts, inter-burst intervals, and total number of sips (Figure S10E-G and S10 K-M). Surprisingly, silencing these neurons did not affect any feeding microstructure parameter for sucrose either (except one driver line for Roundup, leading to a decrease in the length of feeding bursts, Figure S10H-J and S10N-P), suggesting a high level of redundancy within the PER pathway in freely behaving flies. Finally, we decided to silence MN9 (*MN9 SplitGal4* ^19^ > *Kir 2*.*1*) as a final validation of this series of experiments. While there was a clear phenotype in the performance of the underlying motor program (the duration of inter-sip intervals on yeast substrate slightly increased, which is normally not seen under naturalistic conditions) (Figure S11A), the yeast sip durations, sip numbers, inter-burst intervals, and burst lengths were unaffected (Figure S11B-E). However, silencing MN9 led to an increase in sucrose inter-sip intervals as well as the inter-burst intervals (Figure S11F and J), while not affecting sip durations, total number of sips, and burst lengths (Figure S11G-I). This suggests that, similarly to what had been observed in the immobilized PER assays for proboscis extension, MN9 is important for initiating sucrose feeding bursts in freely behaving animals. This finding strengthens our previous results that in freely behaving animals, perturbing neurons in the PER hub does not affect the feeding microstructure underlying yeast consumption. Altogether, we show that CB0499 in the tpGRN PER hub and the previously characterized PER sensorimotor pathway downstream of CB0499 are dispensable for sustaining yeast feeding bursts in yeast-deprived flies. Furthermore, the fact that MN9 silencing increased inter-sip intervals for sucrose but did not affect overall feeding demonstrates that the feeding microstructure in freely behaving animals is highly adaptable and that its underlying circuit is robust to perturbations.

### The tpGRN swallow hub is important for nutrient state-dependent modulation of the burst length

After showing that the PER hub neurons downstream of tpGRNs are dispensable for modulating the length of yeast feeding bursts, we tested whether the tpGRN swallow hub (Figure 4C) is important for controlling the length of feeding bursts. To do so, we created a Split-Gal4 driver line for CB0041 (Figure S12A), which is one of the strongest dtpGRN-specific postsynaptic cell types (Figure S8A). CB0041 (Figure 4H) is a putative glutamatergic neuron (Figure S6) that has high effective connectivity scores with MN10 and MN11D motor neurons of the pharynx, as well as the salivary gland motor neuron MN13^19^ (Figure 4C). Silencing CB0041 led to a slight and almost significant increase in the length of yeast feeding bursts in yeast-deprived flies (p= 0.06, Figure 4J), without affecting the total number of sips and inter-burst intervals (Figure 4I and K). Furthermore, silencing CB0041 in fully fed flies increased the length of yeast feeding bursts significantly without affecting the length of sucrose feeding bursts (Figure S12B and C). CB0041 is a ‘singleton’ cell type consisting of one cell per hemisphere (Figure 4H), which are strongly connected to each other (left-to-right: 59 synapses, right-to-left: 43 synapses, Figure S13A). As glutamate can also be inhibitory in *Drosophila* ^91^, silencing CB0041 could have complex consequences depending on the nature of the glutamate receptors (i.e., excitatory or inhibitory) expressed in CB0041 neurons and their downstream partners. This might explain the increase in the length of feeding bursts upon silencing CB0041. Based on these results, we decided to focus on the sensorimotor pathway downstream of CB0041 to dissect the neural mechanisms underlying nutrient state-dependent modulation of yeast feeding bursts.

### A 3rd-order dtpGRN-specific cell type (Sustain) is important for nutrient state-dependent modulation of yeast feeding bursts

The CB0041 output network (Figure S13A and B) comprises diverse network motifs, including direct and mixed feedback as well as feedforward connections (Figure S13C). To dissect how this network can control the feeding motor program, we first computed the effective connectivity of each of the CB0041 downstream neurons with the proboscis motor neurons. Hierarchical clustering using effective connectivity scores revealed that two of these neurons, CB0874 and CB0302, form a cluster with high effective connectivity scores to motor neurons involved in swallowing: labellar spreading and pharyngeal motor neurons (Figure S14A and B). We focused on these neurons as we showed that the PER pathway is dispensable for modulating the length of yeast feeding bursts (Figure 4, Figure S10 and 11). CB0302 receives 8% (219 of 2678) of its total inputs from CB0041 making CB0041 its second top partner (6% of CB0041’s total outputs) (Figure 5A), while CB0874 receiving only 2.8% (184 of 6490) of its total inputs from CB0041 (5% of CB0041’s total outputs) (for absolute synapse counts see Figure S14C). We therefore hypothesized that CB0302 might be more relevant for controlling the length of yeast feeding bursts, as it is a mutually strong partner with CB0041. Therefore, we turned our attention to dissecting the functional role of CB0302 in burst length modulation. To test if CB0302 is important for modulating the length of yeast feeding bursts, we generated a SplitGal4 driver line targeting CB0302 (Figure S15A-D). A careful analysis of the expression pattern of the line, as well as Multi Color Flip Out (MCFO) ^92^ experiments, revealed that the Split-Gal4 driver line indeed labels CB0302 (Figure 5B and C). We silenced CB0302 using this driver line to test the potential role of CB0302 in modulating the length of yeast feeding bursts. Similar to what we observed upon silencing tpGRNs, CB0302-silenced, yeast-deprived flies could not sustain yeast feeding bursts to the level of the corresponding genetic control (Figure 5D, middle panel). We confirmed the phenotype we observed is due to silencing CB0302 but not the other two cell types labeled by the Split-Gal4 driver line by testing a Split-Gal4 line that labels the other two cell types but not CB0302 (Figure S15E). Indeed, silencing these neurons did not lead to a change in the feeding microstructure (Figure S15G-I). Interestingly, the basal length of yeast feeding bursts remained unaffected upon silencing CB0302 in fully fed flies (Figure 5E, middle panel), and the yeast inter-burst intervals were not affected by CB0302 silencing in either dietary condition (Figure 5D and E, right panels). These findings suggest that CB0302 is necessary to sustain yeast feeding bursts in a nutrient state-dependent way, but dispensable for initiating feeding bursts. We therefore named this neuron “Sustain”. Sustain-silenced flies were able to reach the same amount of total number of sips as the genetic controls within the duration of the experiment (1 hour) in the fully fed condition, and they even had slightly higher total number of sips in the yeast-deprived condition (Figure 5D and E, left panels). These findings demonstrate that flies employ compensatory strategies to reach the same amount of food intake over a longer time range. Next, we tested whether Sustain is necessary for modulating the length of sucrose feeding bursts. Silencing the Sustain neuron in either fully fed or carbohydrate-deprived flies using a carbohydrate-free holidic diet ^93,94^ did not affect the length of sucrose feeding bursts (Figure 5F and G, middle panels). However, we observed a significant decrease in sucrose inter-burst intervals in carbohydrate-deprived flies (Figure 5G, right panel). The total number of sucrose sips was not affected in either condition (Figure 5F and G, left panels). As yeast is the main source of dietary amino acids in flies, we finally asked whether Sustain is important for the modulation of yeast burst lengths upon amino acid deprivation. To test this, we silenced Sustain in amino acid-deprived flies using an amino acid-free holidic diet. Indeed, Sustain was necessary for increasing the length of yeast feeding bursts in amino acid-deprived flies, while being dispensable for modulating the inter-burst intervals and total number of sips (Figure S16). Altogether, we conclude that Sustain is a key neuron that specifically modulates the length of yeast feeding bursts to adapt yeast intake based on the amino acid state of the animal.

**Figure 5.**
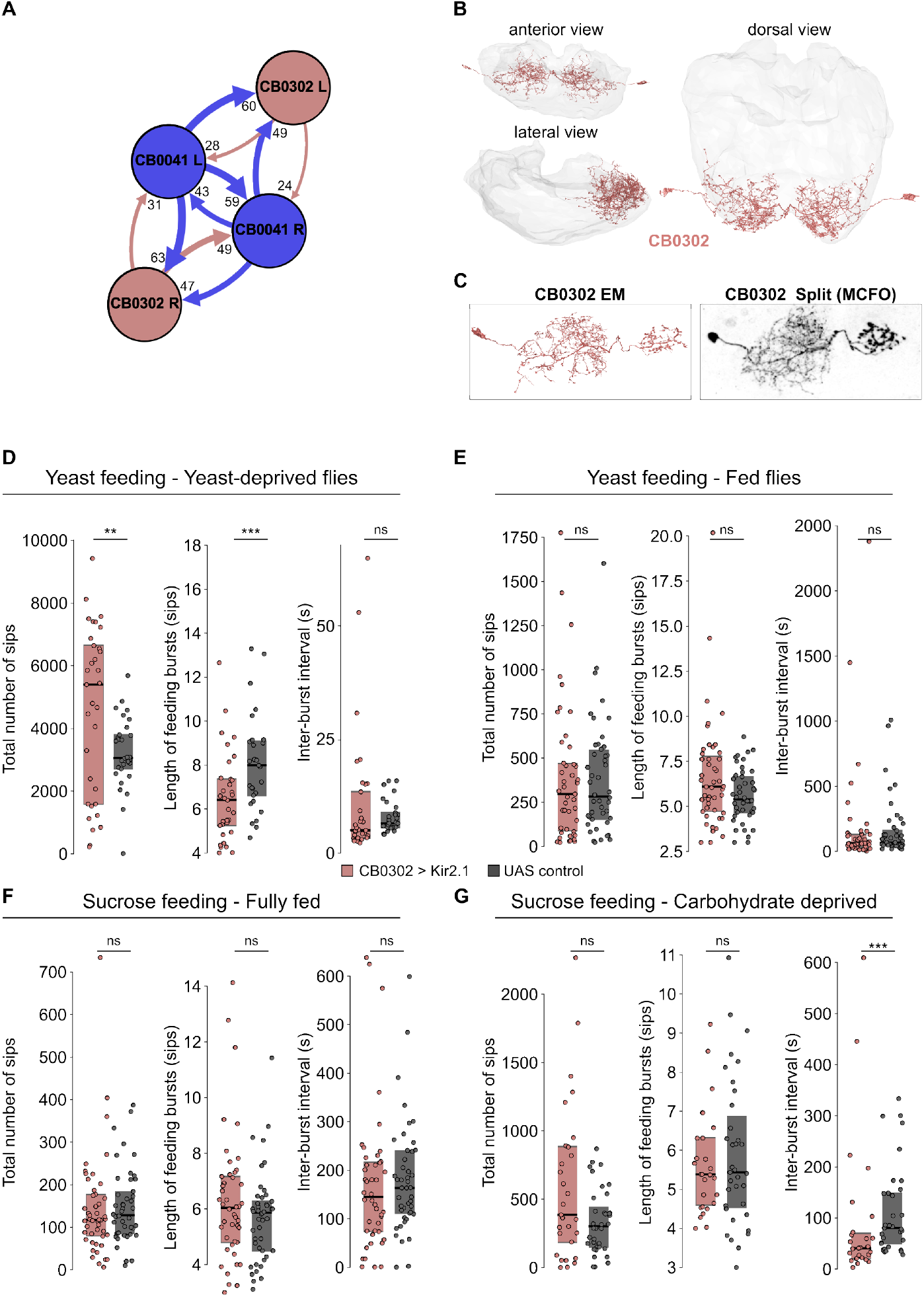
The dorsal tpGRNs pathway neuron CB00302 (Sustain) is important for regulating the length of yeast feeding bursts. **A.** Connectivity diagram between the dtpGRN second-order neuron CB0041 and third-order neuron CB0302. L: left hemisphere, R: right hemisphere. Absolute synaptic weights are shown as numbers beside the arrowheads. **B**. Morphological representation of CB0302 in the EM volume, FAFB – FlyWire. Different points of view are shown. The GNG is shown as a semi-transparent mesh. **C**. Morphological comparison of a CB0302 EM instance (left) with a multi-color-flip-out (MCFO) clone (right) obtained using a specific Split-Gal4 driver line generated to access CB0302. **D**. Yeast-deprived, CB0302-silenced flies have shorter yeast feeding bursts compared to the genetic control (middle panel). They have a higher total number of sips (left panel), but the inter-burst intervals are not affected (right panel). **E**. Fully-fed, CB0302-silenced flies show no difference in the total number of yeast sips, length of yeast feeding bursts, and inter-sip intervals when compared to the genetic control. **F. and G**. Feeding microstructure parameters underlying sucrose feeding are not affected by CB0302 silencing in either fully-fed or carbohydrate-deprived flies. Filled circles indicate individual flies. Boxes indicate the interquartile ranges. Horizontal lines indicate the median value. UAS control: *Empty Split-Gal4* x *UAS-Kir2*.*1*. ns: not statistically significant, Wilcoxon rank-sum test. **: 0.001 *< p <* 0.01, ***: *p <* 0.001.

### Sustain modulates the length of yeast feeding bursts via motor neuron MN11D

To understand how Sustain modulates the length of yeast feeding bursts, we decided to map the downstream circuits that potentially connect Sustain to the actuators of proboscis positioning and ingestion. Sustain has the highest effective connectivity scores with pharyngeal pumping motor neurons (MN11D, 11V, 10, 12D, and MN5 in decreasing order, Figure 6A). Interestingly, after pharyngeal motor neurons labellar spreading motor neurons have the highest effective connectivity score (i.e., MN8 and 7 in decreasing order), which control labellar opening and closing to suck food into the pharynx ^19^. After food is sucked in through the labial palps, pharyngeal motor neurons MN5, 12D, 12V, 11V, and MN11D contract and relax sequentially to pump in the food towards the esophagus ^17,19,41^. Finally, MN10 presumably controls the passage of food from the pharynx to the esophagus, finalizing an ingestion cycle ^19^. Because all these motor neurons underlying ingestion have high effective connectivity scores with Sustain, we hypothesized that modulation of ingestion by the Sustain neuron is important for modulating the length of yeast feeding bursts. As expected, all top downstream partners of Sustain have very high effective connectivity scores with pharyngeal pumping motor neurons except for CB0041, which is mutually connected with Sustain (Figure S17A and B, Figure 5). To test if pharyngeal motor neuron activity is important for modulating the length of yeast feeding bursts, we focused on the pathway to the proboscis motor neuron that has the highest effective connectivity score with Sustain, MN11D (Figure 6A). MN11D receives inputs from Sustain via the premotor neurons CB0910 and CB0459 (Figure 6B, Figure S17C). Since we could not identify a genetic driver line that labels CB0910, we focused on the role of its target, motor neuron MN11D (CB0700, Figure 6C). MN11D is one of the two MN11 types (MN11V and MN11D), which were previously shown to be important for controlling the volume of ingested liquid as well as ingestion rate ^17^. To test if MN11D activity, and therefore ingestion, is involved in modulating the length of feeding bursts, we decided to silence MN11D using a specific Split-Gal4 driver line, which had been previously shown to specifically target that neuron ^19^ (Figure 6D), and quantify the feeding microstructure using the flyPAD system. Silencing MN11D in yeast-deprived flies did not alter the total number of sips on yeast or sucrose (Figure 6E and F, left panels). Similarly, inter-burst intervals were not affected by MN11D silencing (Figure 6E and F, right panels). However, flies with silenced MN11D had shorter feeding bursts for either yeast or sucrose, showing that they fail to correctly control the length of the feeding bursts. These observations suggest that modulation of ingestion is necessary to sustain feeding bursts irrespective of the quality of the food (e.g., protein or carbohydrate), and Sustain, a 3rd order taste peg neuron is necessary for modulating the length of feeding burst specifically for proteinaceous food. It is plausible that parallel and/or redundant pathways might converge on pharyngeal motor neurons to control the length of feeding burst for carbohydrates, and independent pathways control the initiation and sustaining of feeding bursts as probability of feeding initiation could still be modulated upon interfering with ingestion.

**Figure 6.**
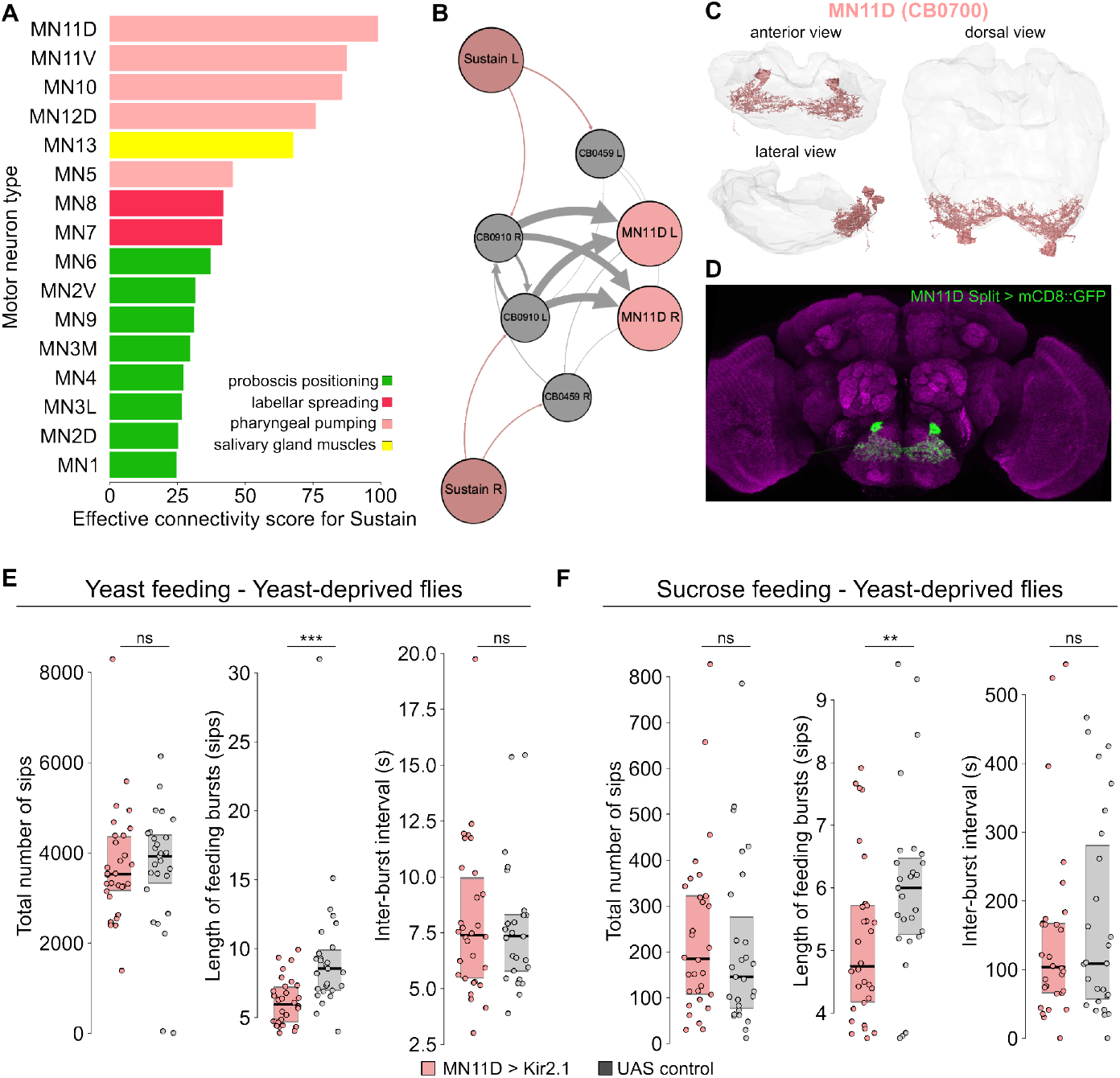
Sustain controls pharyngeal motor neuron MN11D, which is necessary for modulating the length of feeding bursts independent of the nutrient quality. **A**. Effective connectivity scores between Sustain (CB0302) and proboscis motor neurons. Motor neuron types are color-coded based on their function and sorted according to effective connectivity. **B**. Sustain is connected to MN11D through the fourth-order neurons, CB0910 and CB0459. L: left hemisphere, R: right hemisphere. **C**. Morphological representation of MN11D (CB0700) in the EM volume, FAFB – FlyWire. Different points of view are shown. The GNG is shown as a semi-transparent mesh. **D**. The expression pattern of the Split-Gal4 driver line labeling MN11D in the brain (maximum intensity projection). Neuropile: magenta, MN11D > mCD8::GFP: green. **E**. and **F**. Silencing MN11D leads to a decrease in the length of the feeding bursts for both yeast and sucrose feeding (middle panels) while the total number of sips and inter-burst intervals remain unaffected (left and right panels). Filled circles indicate individual flies. Boxes indicate the interquartile ranges. Horizontal lines indicate the median value. UAS control: *Empty Split-Gal4* x *UAS-Kir2*.*1*. ns: not statistically significant, Wilcoxon rank-sum test. **: 0.001 *< p <* 0.01, ***: *p <* 0.001.

### Sustain controls pharyngeal pumping

Our findings prompt the question of whether Sustain modulates ingestion and, through that, the length of yeast feeding bursts. To answer this question, we developed a quantitative pharyngeal pumping assay in which we recorded the cibarium of flies feeding on a 10% yeast solution mixed with a food dye (Erioglaucine 5ug/ml, Materials and Methods) and used DeepLabCut ^95,96^, to track and quantify the dynamics of pumping (Supplemental Video 2). We opted for liquid food delivery using a capillary, as flies feeding on liquid food use their proboscis like a straw to ingest food continuously without retracting the proboscis. This allows us to visualize the cibarium, which would be otherwise obstructed by proboscis extension and retraction during the sipping action of solid food. We tracked multiple points on the labellum and cibarium, as well as the tip of the capillary that delivers the food solution, two bristles on each side of the proboscis (for size normalization purposes across different flies), and a point at the neck position (Figure 7A). Before feeding, the proboscis was typically fully retracted, and the cibarium was not visible (t = -1.3 s, Figure 7A). At feeding initiation, the fly extended its proboscis and opened its labial palps (t = 0 s, Figure 7A). We classified labial opening events by calculating the angle between the three points we tracked on the labellum (left, right, and posterior labial palp, Figure 7A and B). We used the labial opening angle to classify the initiation and termination of a pumping burst (Figure 7D, top panel). We classified the cibarial area using four landmarks surrounding the cibarium (Figure 7C). The obtained area was normalized using the distance between two fixed bristles, which do not move with pumping, to account for slight changes in the head position and inter-individual variability in size (Figure 7C). Shortly after the initiation of feeding (increase in labellum opening angle), the cibarium reaches its maximum area (t = 0.116 s, Figure 7A), indicating that food was pumped into the cibarium. At the end of the pumping cycle, the food was pumped into the esophagus, leaving only a small area and therefore amount of food in the cibarium (t = 0.233 s, Figure 7A). In a pumping burst, the pumping cycle was generally repeated multiple times, as evidenced by the rhythmic change in the cibarial area (Figure 7D, bottom panel). Silencing the Sustain neuron led to a smaller ingested volume of food solution per pumping event when compared to the control background, observable by a shift to smaller values in the kernel density estimates of normalized peak cibarial area (Figure 7E) and the mean peak cibarial area per fly (Figure 7F). However, the power spectral density of ingestion frequency was not significantly different compared to control flies (Figure 7G). These findings are similar to what had been previously reported upon MN11D silencing ^17^, strengthening the finding that MN11D is downstream of Sustain. Although the fly does not perform sipping on liquid food, the peak ingestion frequency is around 5 Hz at 25*°* C (dashed vertical lines, Figure 7G), which corresponds to the proboscis extension and retraction frequency observed during sipping on solid food at this temperature ^25^. This suggests that sipping and ingestion might be temporally coupled. Thermogenetic activation of Sustain neuron at 30*°* C using the temperature-sensitive channel TrpA1 increased the normalized peak cibarial area slightly (Figure S18A and B), without having significant effects on ingestion frequency (Figure S18C). Altogether, these findings demonstrate that Sustain activity is important for controlling ingestion volume per pumping event, which in turn is important for modulating the length of feeding bursts in a nutrient state-dependent way. Based on our findings, we propose a model where the nutrient state-dependent modulation of the tpGRN – Sustain – MN11D pathway leads to more efficient ingestion of food upon gustatory stimulation. This leads to a delay in a feedback mechanism that inhibits sipping action to terminate a feeding burst (Figure 7H and I). Our study demonstrates how low-level motor actions that are controlled by distinct sensorimotor hubs (i.e., proboscis extension/retraction and swallowing) are tightly interlinked via a possible feedback mechanism to modulate the higher-order feeding microstructure based on the physiological needs of the animal.

**Figure 7.**
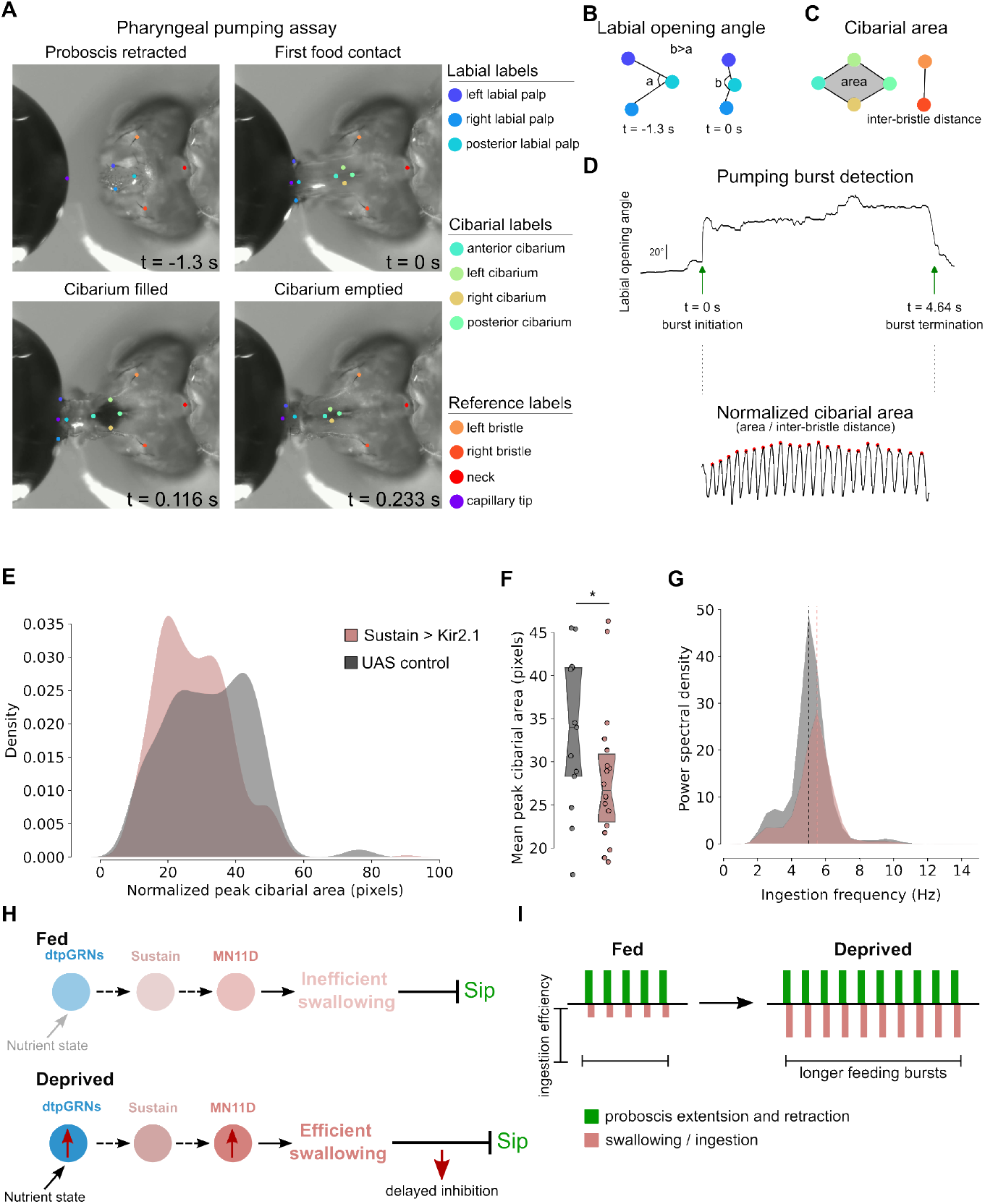
A pharyngeal pumping assay reveals that Sustain modulates the ingestion volume to prolong feeding bursts. **A.** Snapshots of a movie in which a fly was recorded feeding from a capillary filled with artificially colored food (10% yeast with erioglaucine). DeepLabCut was used to track the capillary tip and different points (labels) on the fly to quantify ingestion automatically. The labels are color-coded and annotated on the right side of the panel. Before feeding starts (t = -1.3 s), the proboscis rests in a retracted position. Upon proboscis extension, the proboscis contacts the food droplet, and the labial palps spread to initiate ingestion (t = 0 s, feeding initiation). The fly fills its cibarium (pharyngeal cavity of the fly) with a small amount of food (t = 0.116 s). Finally, the fly empties the cibarium and ingests the food droplet by pumping it through its esophagus (t = 0.223 s). **B**. The labial opening angle is used to identify the initiation of a pumping burst. At rest, the labial angle is small (angle a, t = -1.3 s). Upon contacting the food droplet (t= 0 s), the labial palps spread, leading to an increased labial angle (angle b). **C**. Four points enclosing the cibarium are tracked to calculate the cibarial area. Inter-bristle distances for each fly are used to normalize for the size variation between flies. **D**. Changes in the labial opening angle are used to detect pumping bursts (top panel). A representative time series of normalized cibarial areas (in pixels) during a pumping burst is shown (bottom panel). Red dots represent the peak cibarial area for each swallowing cycle. **E**. and **F**. Sustain silencing leads to a decrease in the ingestion volume as quantified by the normalized peak cibarial area. **E**. Kernel density estimations of normalized peak cibarial areas for Sustain-silenced and control flies. **F**. Mean normalized peak cibarial area for individual Sustain-silenced and control flies. Boxes indicate the interquartile ranges. Horizontal lines indicate the median value. Notches indicate 95% confidence intervals. **G**. Power-spectral density of ingestion frequencies for Sustain-silenced and control flies. Dotted vertical lines represent the peak frequencies. UAS control: *Empty Split-Gal4* x *UAS-Kir2*.*1*. ns: not statistically significant. *: *p <* 0.05. H. and I., the working model. **H**. The modulation of the dtpGRNs pathway by the protein state increases the efficiency of ingestion by anticipatorily activating MN11D before the food arrives in the pharynx, thereby delaying the inhibition of sip generation. **I**. Feeding microstructure model incorporating ingestion. Vertical bars indicate the sipping (green) and ingestion (light pink) actions. The length of the ingestion bars is scaled by efficiency. Because ingestion is more efficient in amino acid-deprived flies, feeding bursts are longer.

## Discussion

Taste-peg gustatory receptor neurons (tpGRNs) are thought to be ideally positioned to evaluate food quality before ingestion: before swallowing, these gustatory neurons have been shown to signal whether a potential meal fulfils the animal’s nutritional needs, especially its demand for amino acids ^39,97^. Here and in an accompanying study ^36^, we combined stateof-the-art EM connectomics, behavioral analysis of feeding, and neurogenetic manipulations to find for the first time that tpGRNs are not a uniform cell type but instead comprise two morphologically and functionally distinct populations: claw tpGRNs (ctpGRNs) and dorsal tpGRNs (dtpGRNs). Using connectivity and morphological clustering alongside computational approaches allowed us to map the connectivity of specific tpGRN types with different proboscis motor neuron (MN) types. We show that these two types feed into separate second-order “hubs”: a proboscis-extension/retraction (PER) hub connected exclusively to ctpGRNs, and a swallowing hub which receives information mainly from dtpGRNs. The swallowing hub second-order neurons preferentially connect to MNs controlling pharyngeal pumping, while the PER hub has high effective connectivity scores with proboscis extension-retraction MNs. Moreover, only dtpGRNs connect to SEZ output neurons, which project to higher brain regions that are involved in navigation, foraging, neurosecretion, learning, and memory, suggesting that dtpGRNs might have further roles in addition to the control of the feeding microstructure ^36^. Together, this dual organization of tpGRN circuitry illustrates how dense connectomic reconstructions and computational approaches can expose hidden cell types and circuit architectures that are almost impossible to identify using classical neuroanatomy and molecular genetics, expanding our understanding of the neural logic that links taste to feeding decisions. This is also the case for other gustatory subpopulations, which we dissect in an accompanying paper ^36^. Building on this anatomical segregation, we first functionally tested the putative PER-arm of the circuit to identify how it could control the feeding microstructure towards yeast. Silencing the principal second-order interneuron Rattle (and each of its downstream neurons: Fdg, Bract, Roundup, MN9) in amino-acid-deprived females had no measurable impact on either the duration or the frequency of yeast-feeding bursts. By contrast, the same manipulations clearly increased the inter-burst interval for sucrose, confirming earlier results that the PER-hub pathway is important for initiating sugar feeding. Given that silencing these neurons did not affect the initiation of yeast feeding bursts, our findings suggest that they are either redundant in freely behaving flies or that there is a parallel pathway mediating the initiation of yeast feeding. In a previous study, we showed that silencing all labial bristle GRNs leads to a decrease in PER response to yeast stimuli, suggesting that labial bristle GRNs are also important for initiating yeast feeding ^39^. Here, we show that IR76b expressing GRNs are important for modulating the frequency of yeast feeding bursts, providing a further indication of the identity of the gustatory bristle neurons promoting yeast feeding initiation and providing an entry point to a future circuit dissection of that pathway. Connectomics analysis of the predicted swallowing hub downstream of dtpGRNs allowed us to identify interneurons that could relay gustatory information coming from this specific subset of the gustatory system. This analysis highlighted a previously unannotated neuron that we termed Sustain. Although not monosynaptically downstream of dtpGRNs, multiple lines of evidence place Sustain at the core of the swallowing hub: it receives strong inputs from dtpGRN-recipient second-order neurons and directs most of its output toward a concise premotor ensemble. Thus, Sustain emerged from the circuit map as a strong candidate for transforming dorsal taste-peg gustatory information into commands that regulate the length of the feeding burst. Furthermore, our circuit analysis predicted, contrary to what we would have hypothesized, that the length of the yeast-feeding burst would be determined by modulating ingestion rather than by the PER control arm. Targeted silencing of Sustain confirmed that this neuron had a nutrient-state-specific role: in aminoacid-deprived females, Sustain inactivation significantly reduced the duration of yeast-feeding bursts without affecting burst frequency or sucrose consumption microstructure. Furthermore, the same manipulation in satiated flies was without effect, indicating that the dtpGRN circuit and therefore also Sustain has a modulatory role whose influence is gated by internal amino acid need rather than acting as a general command for swallowing. These experiments also underscore the value of experimentally perturbing identified neurons to test connectome-based predictions. Indeed, without such manipulations, one might have assumed that the PER circuit, not the swallowing hub, would control feeding-burst length. We show that Sustain is strongly connected to pharyngeal motor neuron MN11D via a set of premotor neurons. Silencing MN11D, which was previously shown to be important for controlling ingestion volume, leads to shorter yeast and sucrose feeding bursts. This finding suggests that ingestion/swallowing efficiency is an important factor controlling the length of feeding bursts independently of nutrient quality. It also suggests that there might be parallel nodes to Sustain controlling sucrose feeding bursts, as Sustain is dispensable for sucrose feeding. We developed a high-speed video assay that quantifies pharyngeal pumping via DeepLabCut-based markerless tracking and showed that Sustain-silenced flies cannot ingest yeast as efficiently as the control flies. This corroborated the model that the dtpGRNs-Sustain-MN11D pathway controls ingestion efficiency for yeast feeding. Based on these findings, we advance a feed-forward facilitation model. When dtpGRNs detect amino-acid–rich food, their activity engages Sustain, which primes MN11D and its premotor partners to increase the volume swallowed in each pump and facilitate cibarium emptying. This boost in swallowing efficacy allows the ongoing proboscis-extension–retraction (PER) cycle to continue seamlessly, producing long, uninterrupted feeding bursts and efficient initiation of the next PER cycle. Conversely, if swallowing cannot keep pace, because MN11D is compromised or Sustain is inactive, the cibarium empties more slowly, bursts terminate early, and subsequent PER cycles stall. Continued sipping would be counterproductive if the previous bolus has not yet been cleared. As the responsiveness of tpGRN to yeast is increased by aminoacid deprivation ^39^, the easiest possible explanation for how internal states modulate the yeast burst length is that Sustain receives stronger dtpGRN-derived input upon aminoacid deprivation, and in turn leads to more efficient swallowing via its connections to MN11D (Figure 7H), ensuring that protein-starved flies maximize ingestion efficiency and, consequently, nutrient uptake. Recently, Ly et al. identified a group of prolactin-releasing hormone (PRLH) neurons in the mouse brainstem, which can control the duration of feeding bursts on time scales similar to the modulation of feeding bursts in flies ^33^. PRLH neurons are necessary for limiting the duration of licking bouts. Unlike Sustain, however, PRLH neurons do not directly control the motor circuits underlying licking, but they rapidly alter the valence of the ingested food. It will be interesting to study if a similar valence-based control of burst length could happen in parallel to the direct motor control of ingestion via Sustain in flies. SEZONs downstream of dtpGRNs are possible candidates to communicate valence as they connect to the mushroom bodies, which are known to encode valence in flies ^98^. Our findings suggest that the ingestion efficiency is a limiting factor for sustaining feeding bursts. How is ingestion efficiency sensed, and how does it modulate the burst lengths? We propose a negative feedback model where pharyngeal sensing of ingested food upon each sip gates the initiation of subsequent sips (Figure 7H and I). Previous studies show that most pharyngeal GRNs mediate aversion, inhibiting feeding ^99^, which our computational analysis of pharyngeal GRN downstream circuits in the accompanying paper also predicts ^36^. Moreover, it was previously suggested that mechanical sensing of water via GR43a+, NOMPC+ sensory neurons in the pharyngeal sense organs limits drinking volume ^72^. Flies could utilize a similar gustatory and/or mechanical sensing mechanism to detect residual food accumulation in the cibarium during a feeding burst to inhibit sipping action. Without the increased activation of the “swallowing hub” circuit upon yeast tasting, food would accumulate more easily in the pharynx, leading to these sensory neurons terminating a feeding burst earlier to avoid mechanical distress. We propose that this sensorimotor pathway should inhibit the motor neurons that control proboscis extension, while promoting swallowing to clean the residual food accumulated in the cibarium and allow the initiation of the next feeding burst. This mechanism could interlink the consequences of two low-level motor actions, proboscis positioning and ingestion, enabling a tight control of the high-level feeding microstructure to control homeostatic feeding. In many animals, including humans, feeding is not a continuous activity. Instead, it takes place in distinct bouts with multiple feeding bursts (meals) of which initiation and duration are tightly controlled. Meal initiation depends on various factors, including circadian rhythm, food availability, and nutrient state ^6,100^. In mammals and flies, it was shown that visceral signals play an important role in determining meal size. In mammals, different segments of the gastrointestinal tract send afferent connections to the brainstem, creating a viscerotopic map in the nucleus of the solitary tract (NTS) ^101,102^. It has been shown that afferents with different G protein-coupled receptor expression profiles respond to nutrient or mechanosensory stimuli. NTS project to behavioral control regions either directly, or indirectly via the lateral parabrachial nucleus (LBPN) ^103^. A recent study demonstrated that distinct neuronal populations in LBPN respond to the passage of food through the gastrointestinal tract, suggesting that food passage is tightly monitored during ingestion and LBPN could signal ongoing food consumption and satiation. In flies, mechanosensation and nutrient sensing in the pharynx and crop were shown to be important for controlling ingestion volume and frequency ^72,104–107^. A recent study showed that enteric GR43a-expressing gustatory receptor neurons sense sugar to gate food passage into the crop based on the nutrient state of the fly ^89^. In addition, nutrient sensing in the gut was shown to control nutrient-specific appetite in flies ^108^. Combined with these findings, our study suggests that monitoring food passage throughout the gastrointestinal tract during ingestion is evolutionarily conserved, and it allows animals to directly control ingestion within timescales compatible with the ingestion cycle and satiation in a feedforward way. We show how recent advances in mapping neural circuits in flies using state-of-the-art EM connectomics are allowing us to understand the neural basis of how external sensing of food is integrated with visceral and humoral signals to control feeding behavior on different time scales ^33^. Furthermore, our study shows how neural systems can orchestrate distinct sensorimotor mechanisms underlying complex and adaptive conserved motor actions that allow animals to attain homeostasis.

## Acknowledgments

We thank Samuel J. Walker for the preliminary dataset that helped us identify the Split-Gal4 driver line targeting Sustain. We thank Julie Simpson for sharing the MN Split-Gal4 driver lines, Gabrielle Sterne and Attilio P. Ceretti for sharing MN annotations in FlyWire, Kathrine Eichler and Isabella Beckett for help with the effective connectivity scripts, Dennis Goldschmidt and the whole Behaviour and Metabolism laboratory for many fruitful discussions, valuable feedback throughout the project, and comments on the manuscript. We thank the Drosophila Connectomics Group, the Jefferis laboratory for fruitful discussions and help with the connectomics analysis. We thank Célia Baltazar and Ana Paula Elias for technical and experimental support. We thank the Champalimaud Fly and Rodent Platforms; the Champalimaud Advanced BioImaging and BioOptics Experimental Platform; the Champalimaud Hardware and Software Platform; the Champalimaud Glass Wash and Media Preparation Platform for help with media preparations. We thank the Princeton FlyWire team and members of the Murthy and Seung labs, as well as members of the Allen Institute for Brain Science, for development and maintenance of FlyWire (supported by BRAIN Initiative grants MH117815 and NS126935 to Murthy and Seung). We also acknowledge members of the Princeton FlyWire team and the FlyWire consortium for neuron proofreading and annotation. Lines obtained from the Bloomington Drosophila Stock Center (NIH P40OD018537) and the VDRC were used in this study. I. T. was supported by a Marie Skłodowska-Curie postdoctoral fellowship (H2020-WF-01-2018-867459), R.B. was supported by a doctoral fellowship 2021.07285.BD from the Portuguese Foundation for Science and Technology (FCT). N. O., G. D. and S. W. were supported by a Wellcome Principal Research Fellowship (200846), a Wellcome Discovery Award (225192), an ERC Advanced Grant (789274), and Wellcome Collaborative Awards (203261 and 209235). The project leading to these results has received funding from FCT (PTDC/MEDNEU/4001/2021), and the “la Caixa” Banking Foundation under the project codes HR17-00539 and HR23-00516 to C.R, Research at the Centre for the Unknown is supported by the Champalimaud Foundation and by Portuguese national funds through FCT - Fundação para a Ciência e a Tecnologia - in the context of the project UIDB/04443/2020, the research infrastructure CONGENTO, co-financed by Lisboa Regional Operational Programme (Lisboa2020), under the PORTUGAL 2020 Partnership Agreement, through the European Regional Development Fund (ERDF) and FCT under the project LISBOA-01-0145-FEDER-022170, and the PPBI - LISBOA-01-0145-FEDER-022122.

## Author contributions

Conceptualization, I.T. and C.R.; methodology, I.T. and C.R.; investigation, I.T., I.V., R.J.B., N.O., and G.D.; formal analysis, I.T., I.V., R.J.B., N.O., and G.D.; visualization, I.T.; writing, I.T. and C.R.; supervision, I.T., S.W., and C.R.; funding acquisition, I.T., S.W., and C.R.

## Resource availability

### Lead contact

Requests for further information and resources should be directed to and will be fulfilled by the lead contact, Carlos Ribeiro.

### Materials availability

This study did not generate new, unique reagents.

### Data and code availability

This paper contains analyses that used existing, publicly available EM connectomics data that can be downloaded (https://codex.flywire.ai/api/download?dataset=fafb). Any additional information required to reanalyze the data reported in this work is available from the lead contact upon request.

## Materials and Methods

### Fly Husbandry

Flies were reared on yeast-based medium (YBM: per liter of water: 8 g agar (NZYTech, Portugal), 80 g barley malt syrup (Próvida, Portugal), 22 g sugar beet syrup (Grafschafter, Germany), 80 g cornflour (Próvida, Portugal), 10 g soya flour (A. Centazi, Portugal), 18 g instant yeast (Safinstant, Lesaffre, France), 8 mL propionic acid (Argos), and 12 mL nipagin (Tegosept, Dutscher, France) (15% in 96% ethanol)) supplemented with instant yeast granules on the surface (Saf-instant, Lesaffre, France) and kept at 22*°* C, 70% relative humidity in a 12 h/ 12 h light-dark cycle. The transgenic fly lines used in this study are shown in Supplementary Table 1.

### Internal state manipulations and holidic media

Holidic media for amino acid and carbohydrate deprivation were prepared as described previously (using the exome-matched formulation with food preservatives described in Piper et al. ^93^). For amino acid-deprived flies, all amino acids contained in the medium were removed from the diet, while keeping the concentration of all other nutrients constant. Likewise, for carbohydrate-deprived flies, all carbohydrates were removed from the medium. For all experiments using the holidic medium, the following dietary treatment protocol was used to ensure a fully fed state: groups of 1 – 4-day-old flies (15 females) were collected into fresh YBM-filled vials, and 5 Canton-S males were added to the vials to ensure females were mated. After 48 hours, flies were transferred to fresh YBM to ensure all flies reach a fully fed state before internal state manipulations. After 24 hours in fresh YBM, flies were transferred to different holidic media and kept there for 3 days. In yeast deprivation experiments, yeast deprivation was induced by transferring 2-6 day-old, mated and fully fed female flies to vials containing a paper tissue soaked with 5 mL of 100 mM sucrose solution for 5 days. Fully fed flies were age-matched with protein-deprived flies. Mating was ensured by keeping 15 female flies together with 5 male flies.

### flyPAD assays

The feeding microstructure was monitored using flyPAD as described in Itskov et al ^25^. Female flies with desired genotypes were placed in an arena with two electrode pairs (channels) filled with food patches containing 1% agarose mixed with either 10% yeast or 20 mM sucrose. The flies were allowed to feed for one hour, and sips on either food patch were extracted from the capacitance traces read from either channel using previously described algorithms. Briefly, capacitance traces were acquired using the Bonsai framework (versions 2.1.4 – 2-8.5) at 100 Hz and analyzed in MATLAB (version 8.2 (R2013b)) and Python (3.9.15) using custom-written software. After extracting sip forms, feeding bursts and activity bouts were identified by threshold-based criteria as described previously. The minimum length of a feeding burst is defined as three sips. Non-eating flies (those having less than two activity bouts per assay) were excluded from the analysis. flyPAD experiments were performed at 25° C with 70% relative humidity.

### Manipulation of neuronal activity

To silence distinct neuronal cell types, Kir2.1 was expressed in different neuronal subsets using Split-Gal4 drivers (using a *5X-UAS:Kir2*.*1* transgene). To induce the expression of Kir2.1, flies were moved to 30*°* C 24 hours before the behavioral experiments. For thermogenetic activation, TrpA1 was expressed in the Sustain neuron using a specific Split-Gal4 driver (using a *UAS-trpA1* transgene). Thermogenetic activation experiments were performed at 30*°* C. The details of the Split-Gal4 driver lines used in this study are shown in Supplementary Table 1.

### Immunostaining

Brains of 8-10 days old, fully fed, mated females expressing mCD8:GFP under the control of specific Split-Gal4 driver lines were used (using a 20X-UAS-mCD8:GFP transgene). Brains were dissected in 4° C 1X PBS (10173433, Fisher Scientific, UK) after a quick passage through EtOH (4146052, Carlo Elba) and were then transferred to formaldehyde solution (4% paraformaldehyde, P6148, Sigma-Aldrich in 1X PBS + 10% Triton-X, X100, Sigma-Aldrich) and incubated for 20-30 min at room temperature. Samples were then washed three times in PBST (0.5% Triton-X in PBS) and then blocked with 10% normal goat serum (16210-064, Invitrogen) in PBST for 15-60 min at room temperature. Samples were then incubated in primary antibody solutions (Rabbit anti-GFP, TP401, Torrey Pines Biolabs at 1:6000 and Mouse anti-NC82, Developmental Studies Hybridoma Bank at 1:10 in 5% normal goat serum in PBST). Primary antibody incubations were performed for 3 days at 4° C with rocking. They were then washed in PBST 2-3 times for 10-15 min at room temperature. The secondary antibodies were applied (Anti-mouse A594, A11032, Invitrogen at 1:500 and Anti-rabbit A488, A11008, Invitrogen at 1:500 in 5% normal goat serum in PBST) and brains were then incubated for 3 days at 4° C with rocking. They were again washed in PBST 2-3 times for 10-15 min at room temperature. Samples were mounted in Vectashield Mounting Medium (H-1000, Vector Laboratories). For MultiColorFlipOut experiments, Bloomington stock BDSC#64086 (MCFO-2) was crossed to the Gal4 lines. Adult, 2-4 days old flies were heat shocked for 12-20 minutes at 37° C. Heat-shocked flies were kept at 25° C for 3-5 days before dissection. For MultiColorFlipOut immunostainings, the Janelia FlyLight protocol (IHC-MCFO, https://www.janelia.org/project-team/flylight/protocols) was used with a slight modification: the dehydration, clearing, and DPX embedding steps were replaced with mounting in Vectashield Mounting Medium (VectorLabs, USA). Images were captured on an inverted Zeiss LSM 980 (Carl Zeiss Co., Oberkochen, Germany) using a Plan-ApoChromat 20x/0.8 air lens objective (Carl Zeiss Co., Oberkochen, Germany).

### Connectome dataset

Taste peg gustatory receptor neuron projections were identified in the FAFB – FlyWire EM volume (https://flywire.ai/) based on their characteristic morphological features. Incomplete projections were proofread and corrected in the production dataset using the dedicated neuroglancer interface (https://ngl.flywire.ai/). The body IDs of taste peg GRNs are shown in Supplementary Table 2. Proofreading is a community effort as described in Dorkenwald et. al ^44^. FlyWire Brain Dataset version 783 (https://codex.flywire.ai/) was used for the connectomics analysis in this study. Synapse detection was based on Buhmann et al ^43^. Neurotransmitter predictions were based on Eckstein et al ^47^. Cell type annotations by Schlegel et al. ^46^ were used in this study.

### Connectome analysis

Natverse package ^109^ in R (R Development Core Team, version 4.1.0) was used to analyse the connectome dataset. The dplyr package (https://github.com/tidyverse/dplyr) was used to facilitate data manipulation. To retrieve the synaptic-level connectome dataset (FlyWire, v783) and plot meshes, the fafbseg package (urlhttps://github.com/natverse/fafbseg) was used. For synaptic connectivity predictions, we used a cleft threshold = 50. Synaptic weight thresholds were indicated whenever required. Synaptic partners that have less than two synapses were excluded from the analysis. The NBLAST method was used to perform morphological comparison of neuronal anatomies ^79^. The nhclust function in the nat.nblast package was used to hierarchically cluster neurons based on NBLAST scores. Cosine similarity scores were computed using the coconatfly package (https://github.com/natverse/coconatfly). Briefly, this method computes the cosine similarity for pairs of connectivity vectors for neurons in the n-dimensional synaptic connectivity space. Average silhouette width for obtaining the optimal number of clusters was performed using the fviz_nbclust function in the factoextra package (https://github.com/kassambara/factoextra). Graph representations of neuronal connectivity were plotted using Gephi (https://gephi.org/, version 0.10) to highlight the relative connectivity of nodes. The neurons were treated as nodes, and the synaptic connectivity between neurons was treated as directed edges scaled by the synaptic weights for each neuron pair. For the layout of the graphs, the Built-in YiFan Hu Proportional method in Gephi was used. YiFan Hu’s method is a time-efficient method based on force-directed algorithm models that aim to minimize the energy in the system iteratively. The effective connectivity algorithm was adopted from Sturner et al ^87^, as explained in detail in the companion paper ^36^.

### Pharyngeal pumping assay

Female flies with desired genotypes were mounted by gluing the thorax and abdomen dorsally on a coverslip. Flies were acclimatized for 30 mins at 25° C, 70% humidity before the initiation of the experiment. The coverslip was flipped and mounted on a holder. A high-performance camera (Grasshopper 3 – USB3, Teledyne Vision Solutions, USA) and a macro lens (Laowa 25mm, f/2.8, 2.5-5X Ultra Macro, Venus Optics, China) were used to visualize the proboscis and cibarium with a ventral view. The videos (H264 – MP4, 2048 x 1536 pixels) were recorded at 60 Hz using Spinnaker SDK (Teledyne Vision Solutions, USA). NanoJect III (Drummond Scientific Company, USA) was used to deliver small drops of dyed food via a capillary (Wiretroll II, 5-000-2005, Drummond Scientific Company, USA). The food was prepared by mixing 10% yeast (Saf-instant, Lesaffre, France) with 5mg/mL Erioglaucine (861146 – 25g, Sigma) in Mili-Q water (Merck, Germany).

### Pharyngeal pumping analysis

Pharyngeal pumping videos were analyzed using DeepLabCut version 2.2.298. The points to be tracked were labeled using the graphical user interface. The network was trained for 758.000 iterations. Likelihood thresholds were used to filter the data points for each tracked point upon careful visual inspection. Custom-written Python (3.9.15) scripts were used to analyze the data. Pumping bursts were detected by calculating and thresholding the changes in the labial opening angle. The cibarial area was calculated based on the four points (anterior, posterior, left, and right) around the cibarium using basic trigonometric functions in the Python package NumPy (v2.1). The signal was filtered with a median filter using the medfilt function (kernel size= 3) in SciPy (v1.15). Only the first pumping bursts for each fly were analyzed. For power spectral analysis, Welch’s method (scipy.signal.welch in SciPy package v1.15) was used with a window size of 120 frames (overlap = 60 frames) after filtering the data with a bandpass Butterworth Filter between 2 and 10 Hz (order = 7, scipy.signal.butter SciPy package v1.15). Peak detection was performed using the find_peaks function in SciPy (v1.15).

### Statistical Analysis

Statistical comparisons for flyPAD data were performed using the Wilcoxon rank-sum test. The ranksums test in SciPy (v1.15) was used. Whenever multiple comparisons were needed, the p-value was adjusted using the Bonferroni correction. Kernel density estimates were computed using the kdeplot function in the seaborn package (v0.13). Scott’s rule was used as the bandwidth method. A bandwidth value of 1 is used. Notches in the notched boxplots indicate 95% confidence interval upon bootstrapping times.

