## Supplemental figures for "Connectomics Reveals a Feed-Forward Swallowing Circuit Driving Protein Appetite"

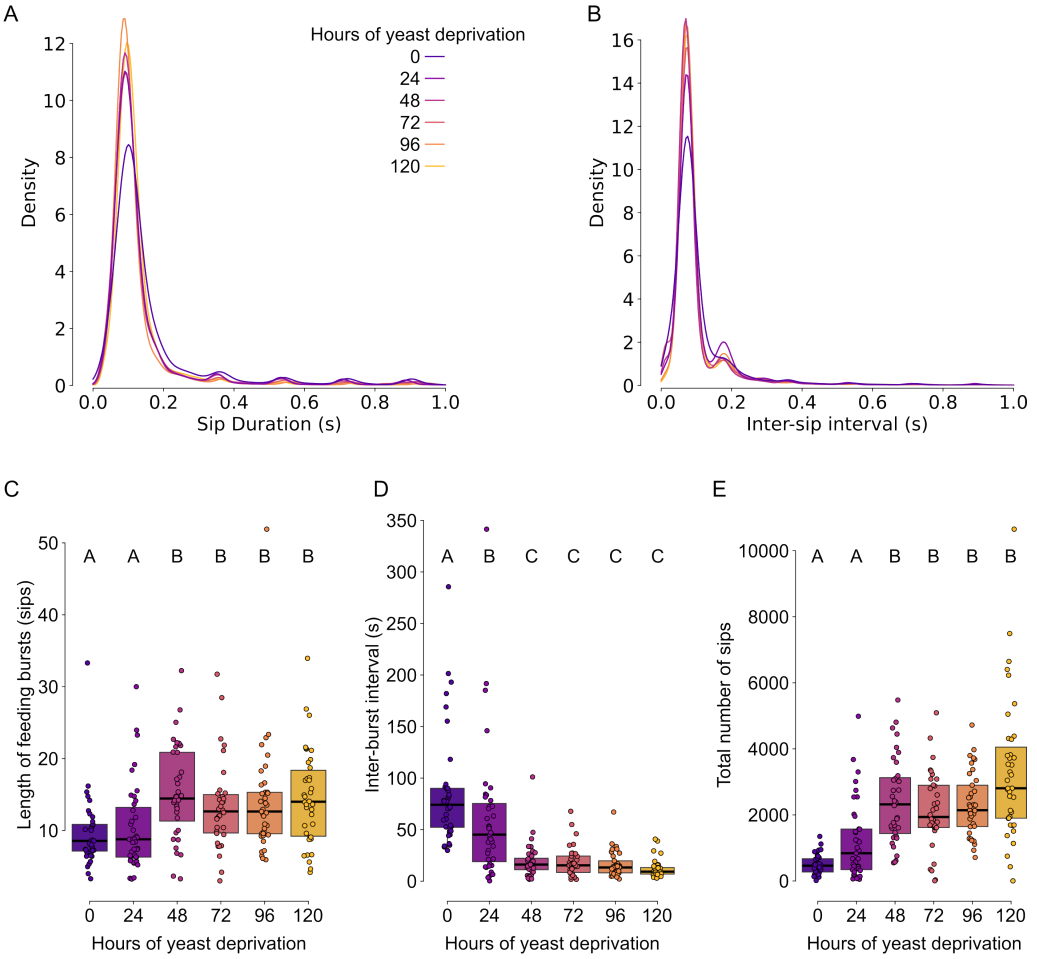


**Figure S1: Modulation of the feeding parameters upon different durations of yeast deprivation. A.** Kernel density estimations for yeast sip durations and **B.** inter-sip intervals for flies deprived of yeast for different durations (Fully fed = 0 hours). **C-E.** Changes in the yeast feeding microstructure parameters upon different durations of yeast deprivation. Different letters indicate statistically significant differences. Filled circles indicate individual flies. Boxes indicate the interquartile ranges. Horizontal black lines indicate the median value. Canton-S flies were used for this experiment.

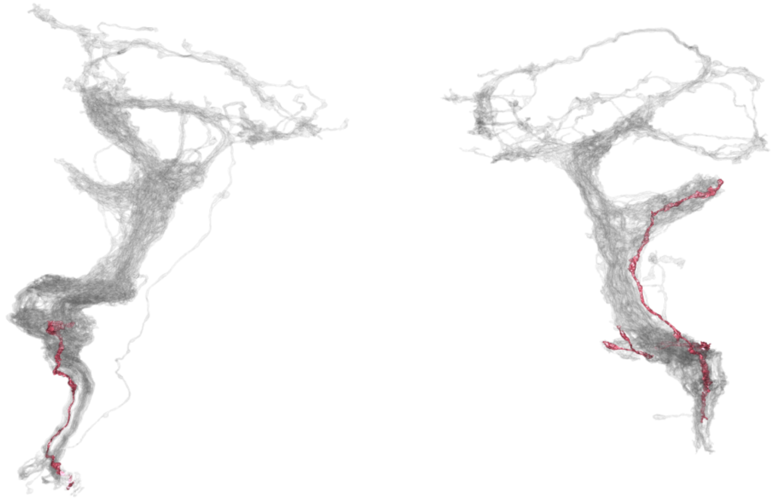

**Figure S2: Incomplete taste peg GRNs.** Incomplete taste peg GRNs in the EM volume, FAFB – FlyWire. Fully proofread tpGRNs are shown as semitransparent grey mesh. Anterior view.

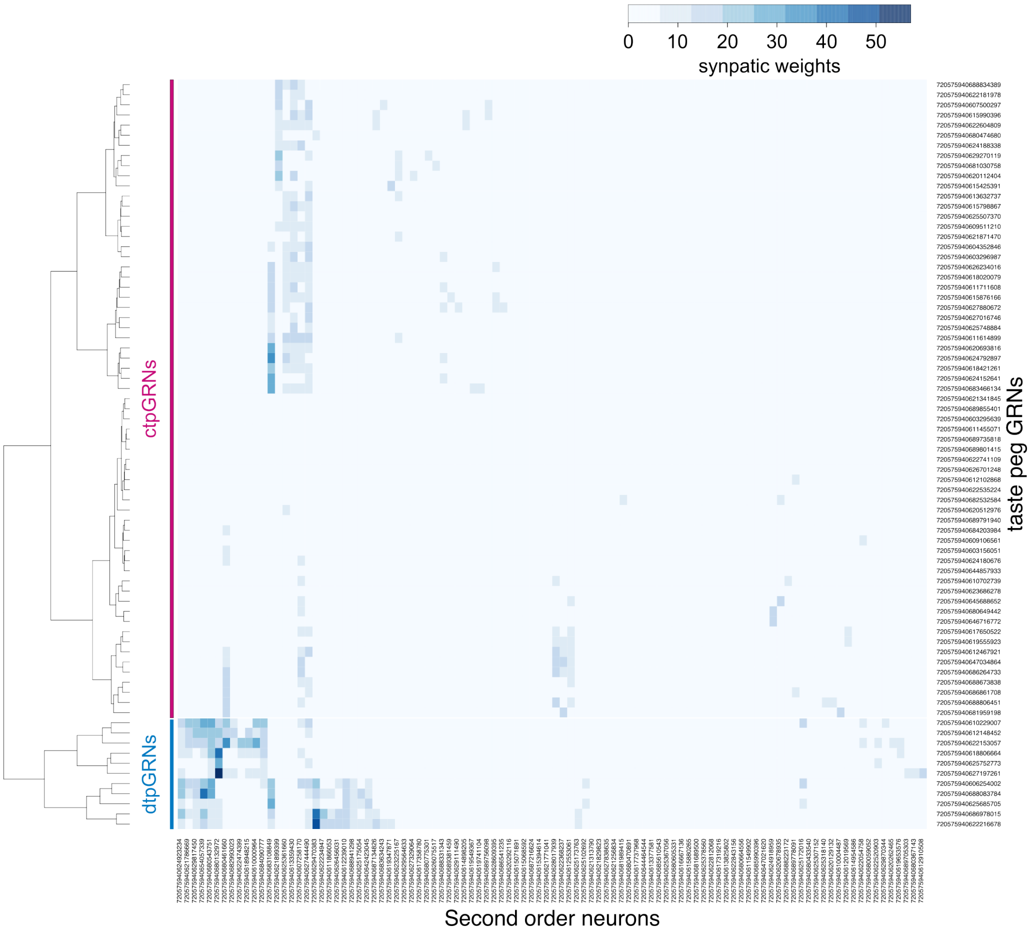


**Figure S3: tpGRN downstream connectivity and functional tpGRN type.** tpGRN downstream connectivity is shown as a heatmap. The FlyWire segment IDs (version 783) are indicated for both tpGRNs and second-order neurons. The synaptic strength between each tpGRN–second-order neuron pair is color-coded. Hierarchical clustering (Ward's method) using synaptic strengths revealed the dtpGRN (blue) and ctpGRN (magenta) as two functionally distinct tpGRN subtypes.

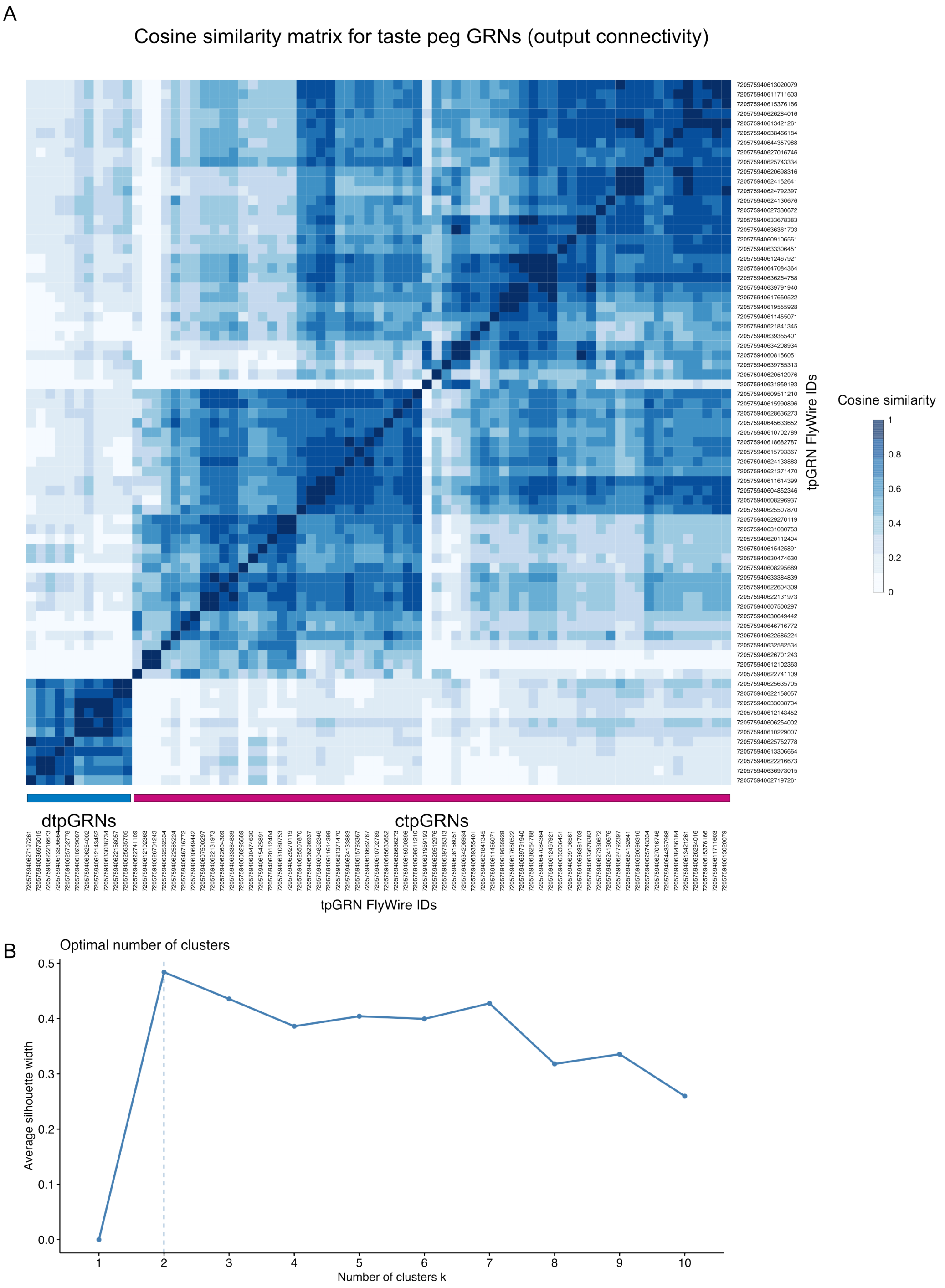


**Figure S4: Cosine similarity of dtpGRNs and functional tpGRN types. A.** Cosine similarity matrix of tpGRNs is shown as a heatmap. Cosine similarity (0-1) is color-coded. The FlyWire segment IDs (version 783) are indicated for both tpGRNs and second-order neurons. Hierarchical clustering (Ward's method) using cosine similarity of tpGRNs revealed the dtpGRNs (blue) and ctpGRNs (magenta) as two functionally distinct tpGRN subtypes. **B.** Average silhouette width analysis based on the cosine similarity matrix suggests there are optimally two tpGRN clusters.

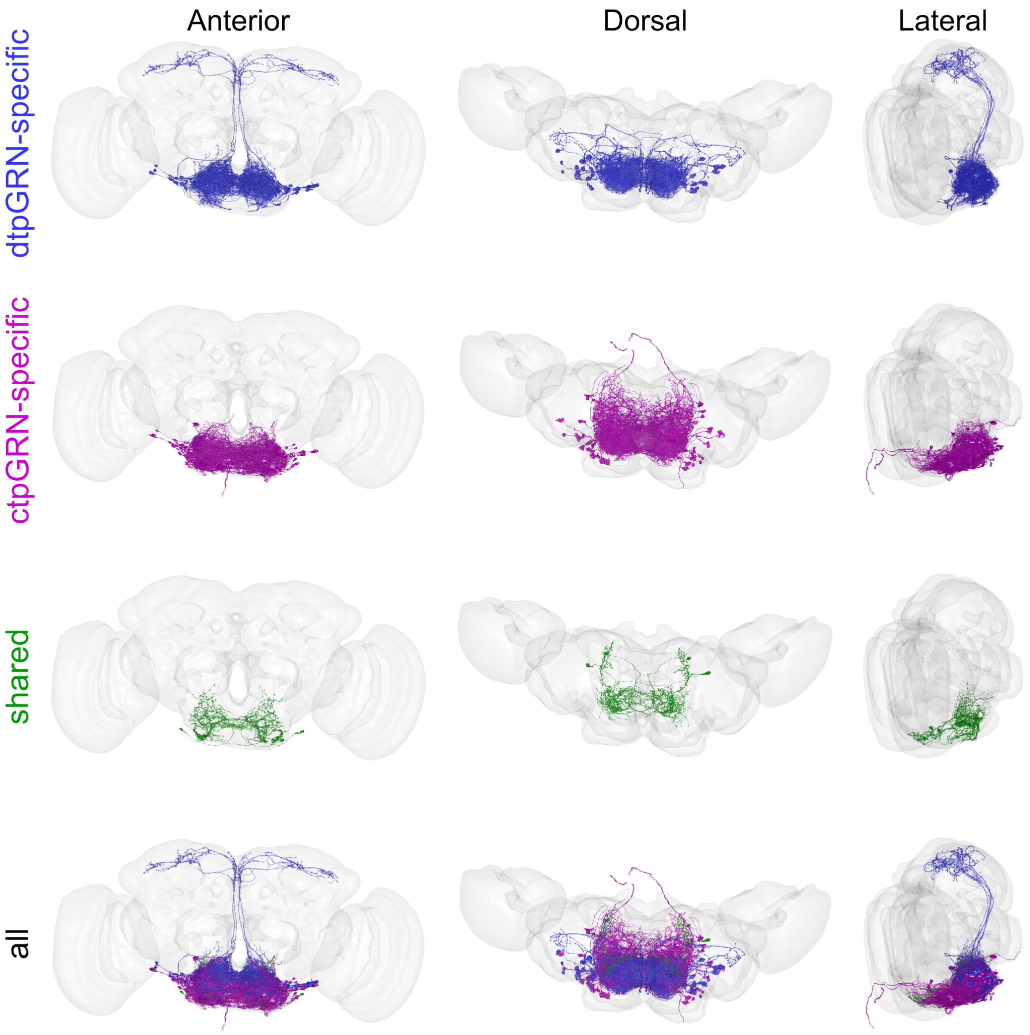


**Figure S5: Anatomy of tpGRN downstream partners.** Morphological representation of tpGRN downstream partners in the EM volume, FAFB – FlyWire. Different points of view (anterior, dorsal, and lateral) are shown. The neuropile is shown as a semi-transparent mesh. Blue (top row): dtpGRN-specific downstream partners, magenta (second row): ctpGRN-specific downstream partners, green (third row): shared downstream partners, overlay (bottom row).

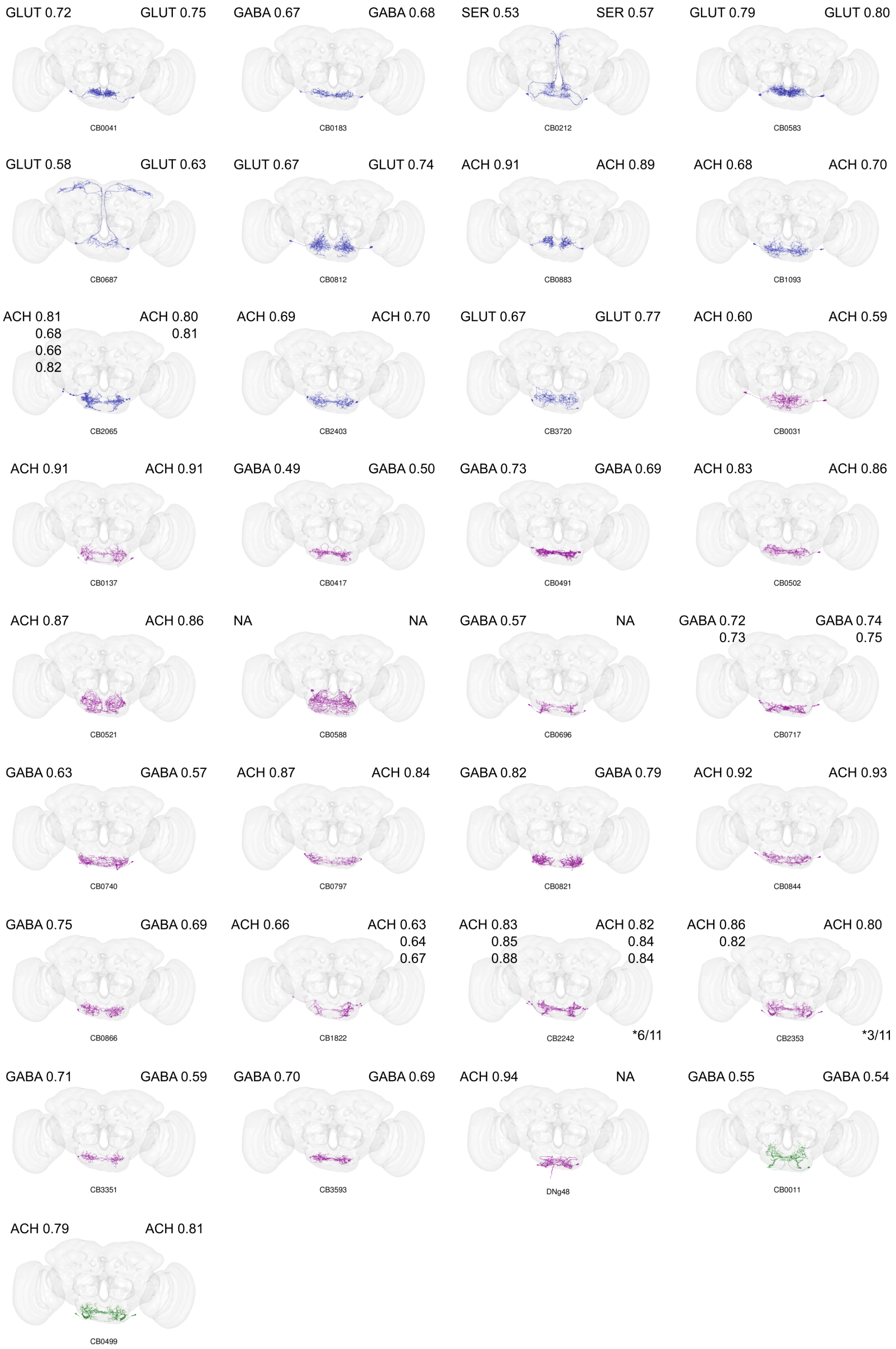


**Figure S6: Anatomy of each tpGRN downstream partner cell type and neurotransmitter predictions.** Morphological representations of tpGRN downstream cell types with the top neurotransmitter predictions for each cell in either hemisphere. Some cell types have more than two cells per hemisphere. Neurotransmitter predictions per cell are shown. Each cell type is color-coded according to whether they are dtpGRN-specific (blue), ctpGRN-specific, or shared (green). ACH: acetylcholine, GABA: gamma-aminobutyric acid, GLUT: glutamate, SER: serotonin, NA: no prediction.

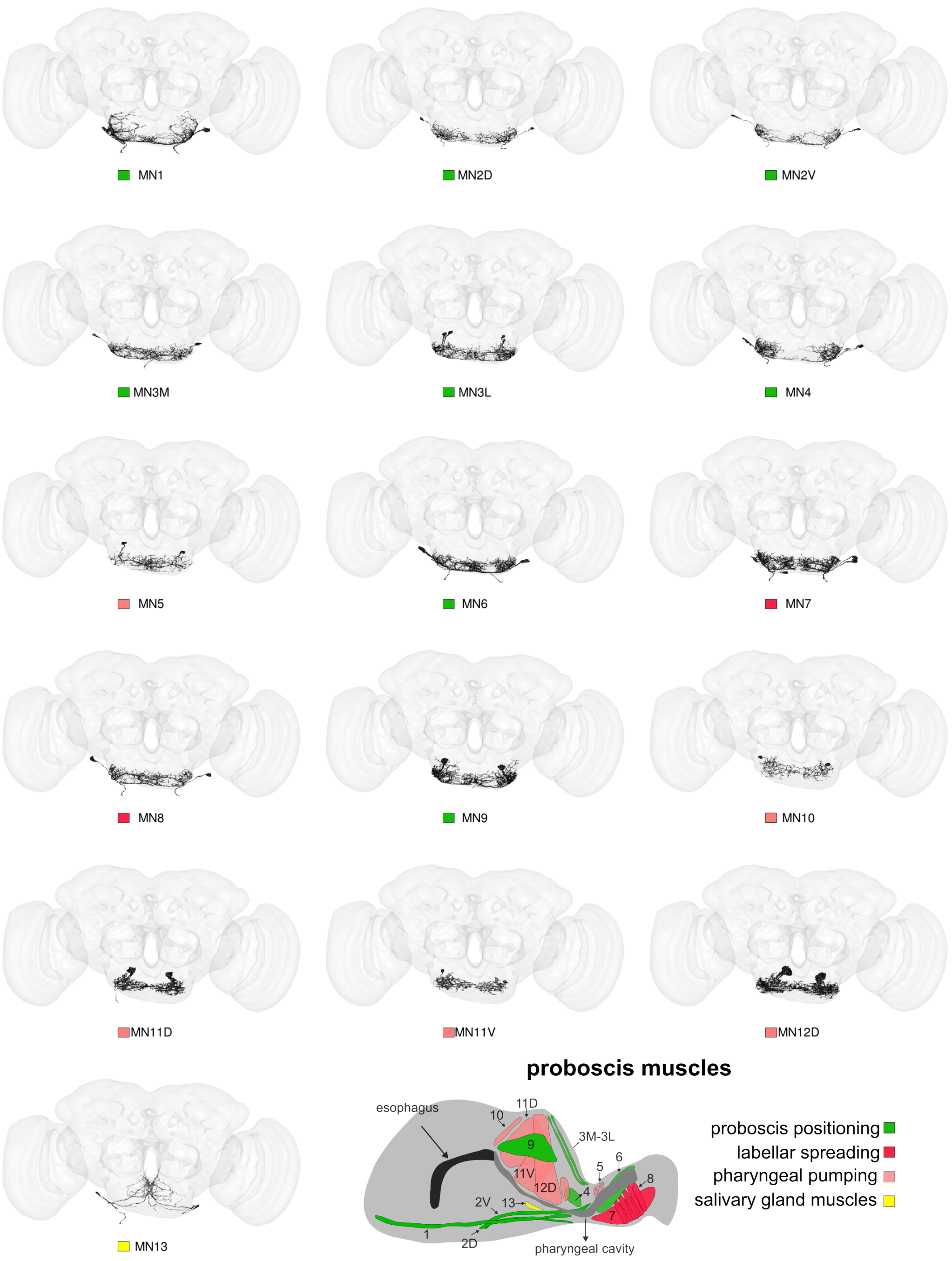


**Figure S7: Anatomy of proboscis motor neurons and their functions.** Morphological representations of proboscis motor neuron instances identified in the FAFB – FlyWire volume. The colored rectangle for each motor neuron indicates the function that the motor neuron is involved in, as shown in the schematic (right, bottom row).

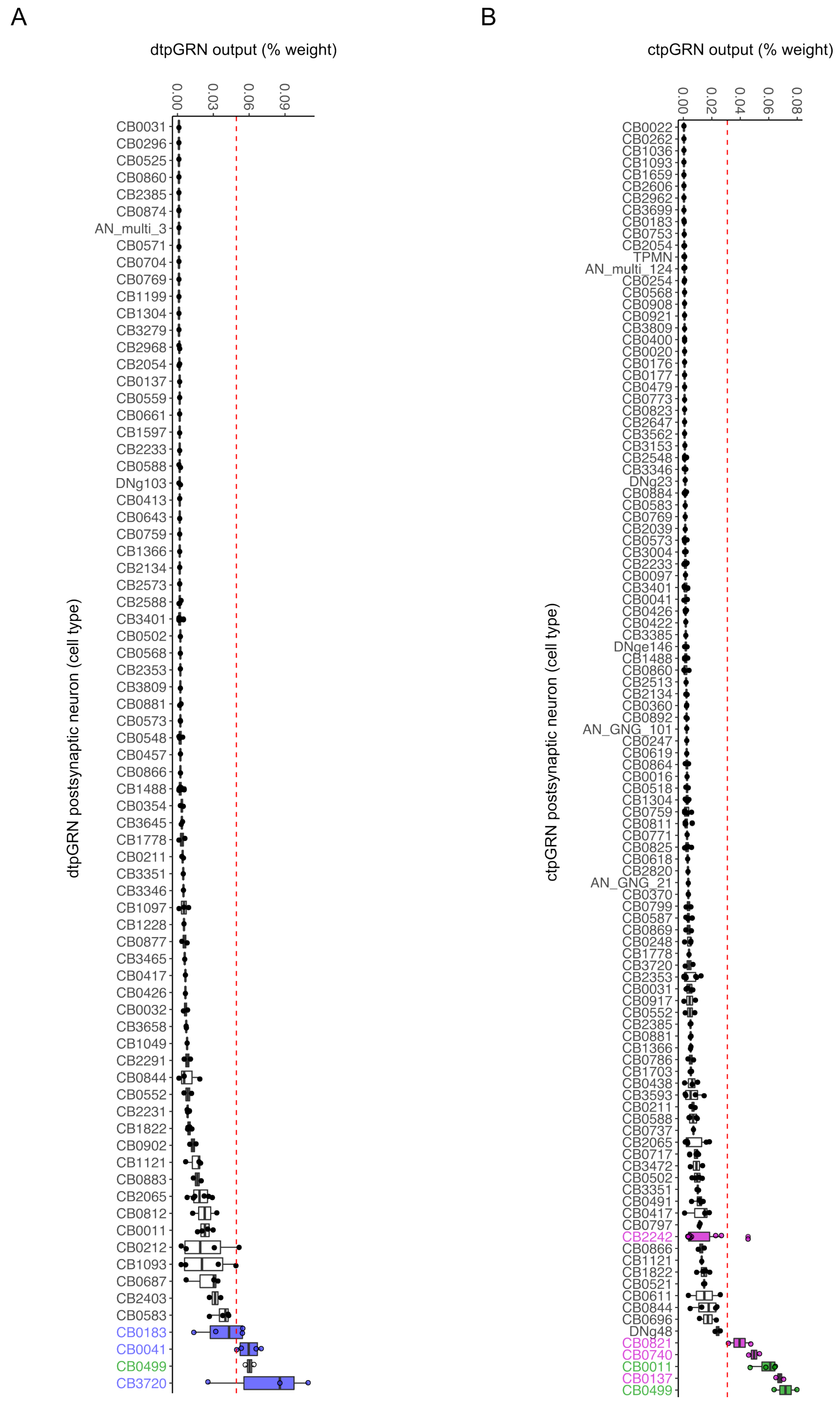


**Figure S8: Proportional outputs of dtpGRNs and ctpGRNs onto their downstream cell types. A.** Proportional connectivity between dtpGRNs and all dtpGRN downstream partners (proportion of total dtpGRN outputs). **B.** Proportional connectivity between ctpGRNs and all ctpGRN downstream partners (proportion of total ctpGRN outputs). Blue: top dtpGRN-specific, magenta: top ctpGRN-specific, green: top shared downstream partners. Each filled circle represents a single downstream neuron. Boxes indicate the interquartile ranges. Vertical black lines indicate the median value. Vertical dashed lines (red) indicate the 95th percentile for the distribution of synaptic weights for all downstream partners for **A.** dtpGRNs and **B.** ctpGRNs. If a cell type has at least two neurons with synaptic weights bigger than the 95th percentile, it is considered a top partner.

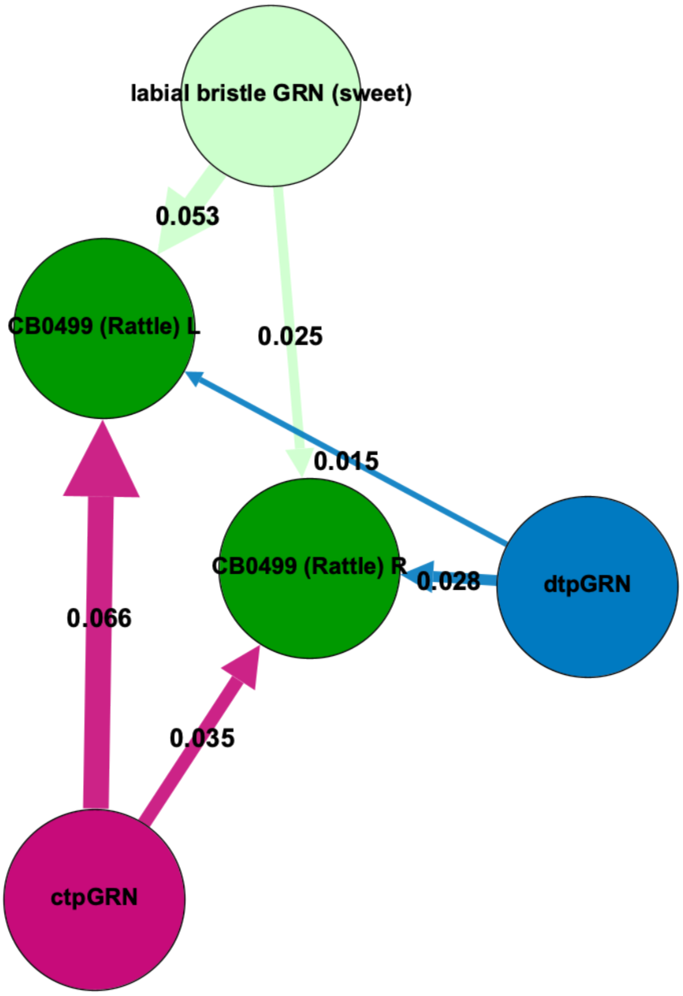


**Figure S9: GRN inputs of CB0499.** Upstream connectivity of the CB0499 (Rattle) neuron. The proportional synaptic inputs to Rattle from tpGRN types and sweet-sensing labial bristle gustatory receptor neurons are indicated over the edges. L: left, R: right.

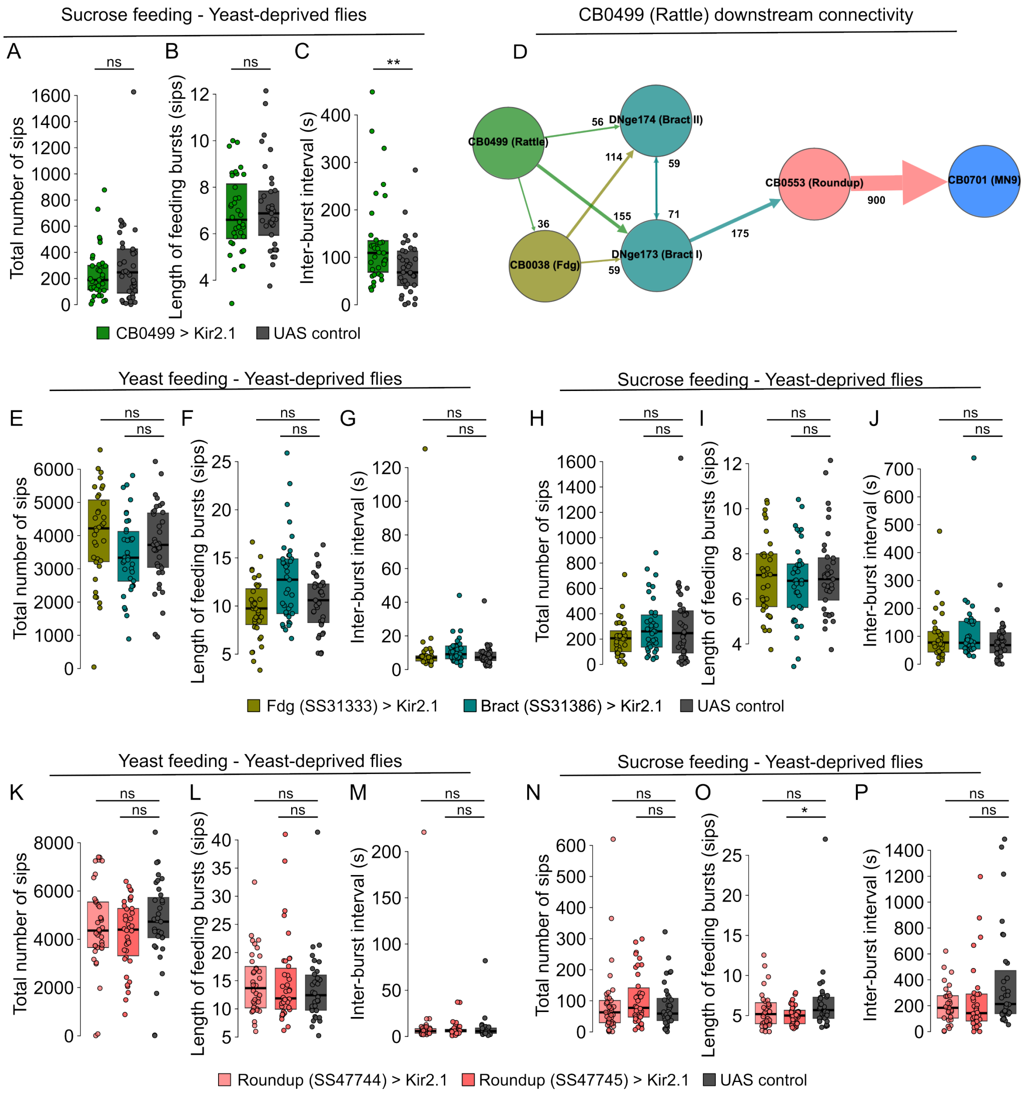


**Figure S10: Loss-of-function experiments for the PER pathway. A-C.** Silencing CB0499 in yeast-deprived flies leads to longer inter-burst intervals while not affecting the total number of sucrose sips or the length of sucrose feeding bursts. **A.** Total number of sips. **B.** Length of feeding bursts. **C.** Inter-burst intervals. **D.** The cell types connecting CB0499 to MN9. Absolute synaptic weights between cell types are indicated over the edges. **E-J.** Silencing Fdg or Bract using Split-Gal4 driver lines SS31333 and SS31386 did not affect any feeding parameters on either yeast or sucrose. **K-P.** Silencing Roundup using two different Split-Gal4 lines (SS47744 and SS47745) did not affect any feeding parameter, with the only exception that silencing SS47745 leads to a slight decrease in the length of sucrose feeding bursts. UAS control: *Empty Split-Gal4* x *UAS-Kir2.1*. Filled circles indicate individual flies. Boxes indicate the interquartile ranges. Black horizontal lines indicate the median value. ns: not statistically significant, Wilcoxon rank-sum test. *: $0.01 < p < 0.05$, **: $0.001 < p < 0.01$.

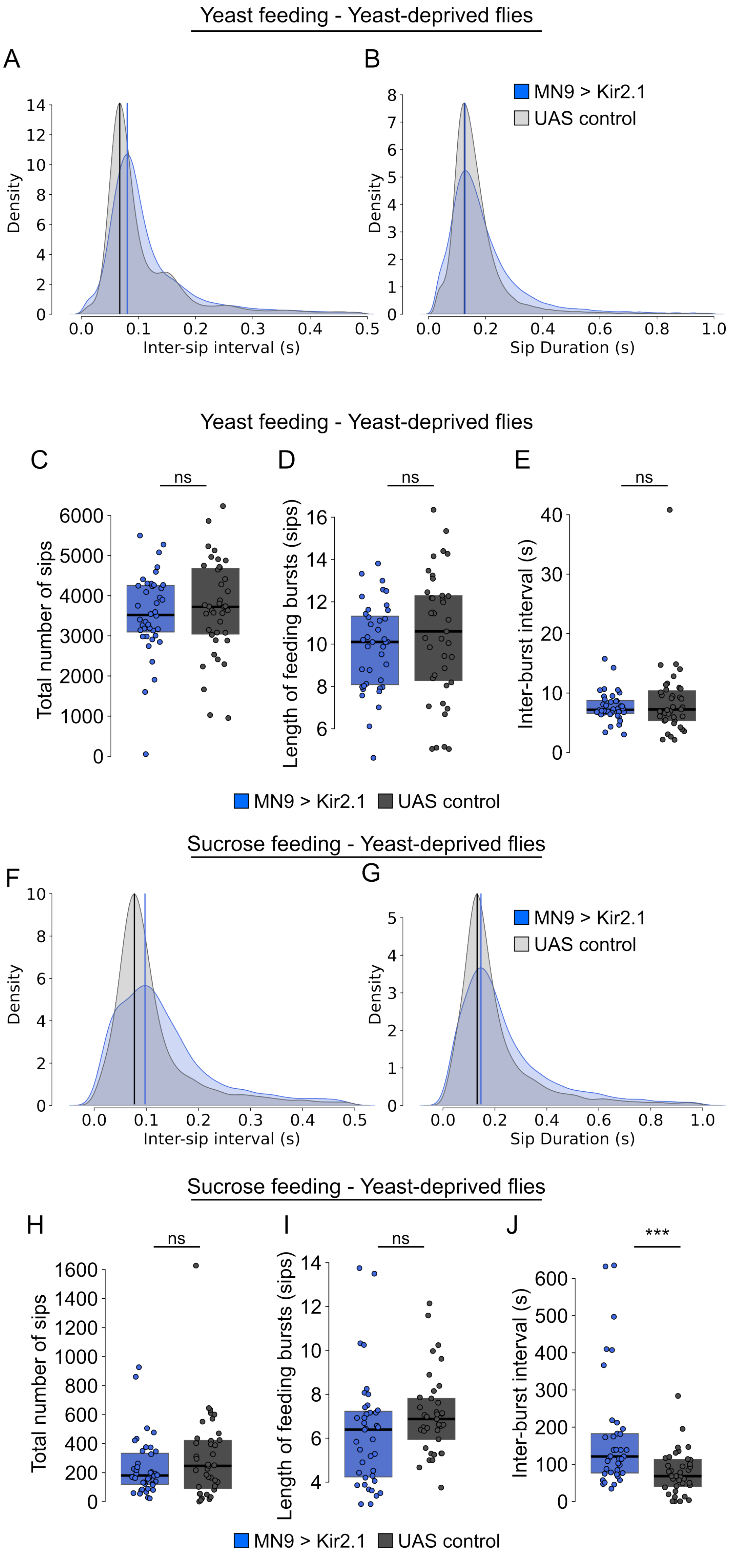


**Figure S11: Effects of MN9 silencing on yeast and sucrose feeding.** Silencing MN9 (blue) leads to a slight shift in the kernel density estimates for **A.** inter-sip intervals compared to the controls (grey), while not affecting **B.** sip durations for yeast feeding. Vertical lines indicate peak values for inter-sip intervals and sip durations. **C-E.** Silencing MN9 does not affect the **C.** total number of sips, **D.** length of feeding bursts, or **E.** inter-sip intervals for yeast feeding. Silencing MN9 (blue) leads to a slight shift in the kernel density estimates for **F.** inter-sip intervals, compared to the controls (grey), and a smaller shift in **G.** sip durations for sucrose feeding. Vertical lines indicate peak values for inter-sip intervals and sip durations. **H-J.** Silencing MN9 does not affect the **H.** total number of sucrose sips or **I.** length of sucrose feeding bursts, but **J.** leads to longer inter-burst intervals for sugar feeding. UAS control: *Empty Split-Gal4* x *UAS-Kir2.1*. Filled circles indicate individual flies. Boxes indicate the interquartile ranges. Horizontal lines indicate the median value. ns: not statistically significant, Wilcoxon rank-sum test. ***: $p < 0.001$.

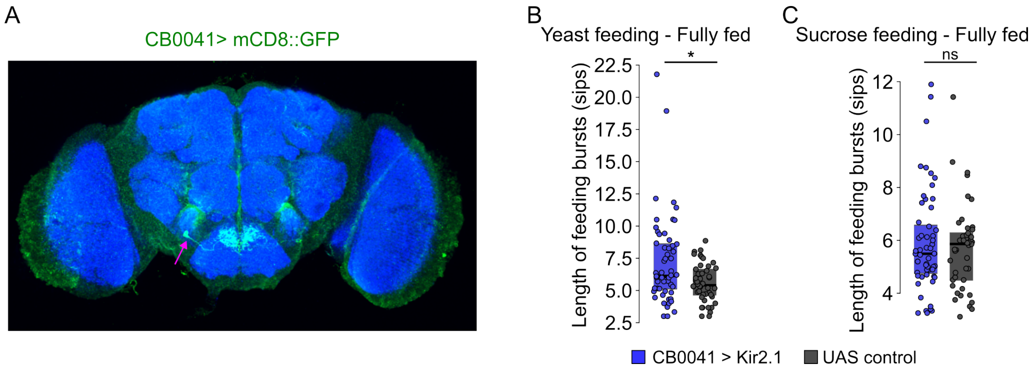


**Figure S12: Characterization of the Split-Gal4 driver line labeling CB0041 and the effect of silencing CB0041 in fully-fed flies. A.** Expression pattern of a Split-Gal4 driver line labeling CB0041 (maximum intensity projection). green: CB0041 neurons expressing mCD8::GFP, blue: neuropile staining (nc82). The arrow indicates the cell body. **B.** Silencing CB0041 in fully-fed flies leads to longer yeast feeding bursts while not affecting **C.** the length of sucrose feeding bursts. UAS control: *Empty Split-Gal4* x *UAS-Kir2.1*. Filled circles indicate individual flies. Boxes indicate the interquartile ranges. Horizontal lines indicate the median value. ns: not statistically significant, Wilcoxon rank-sum test. *: $0.01 < p < 0.05$.

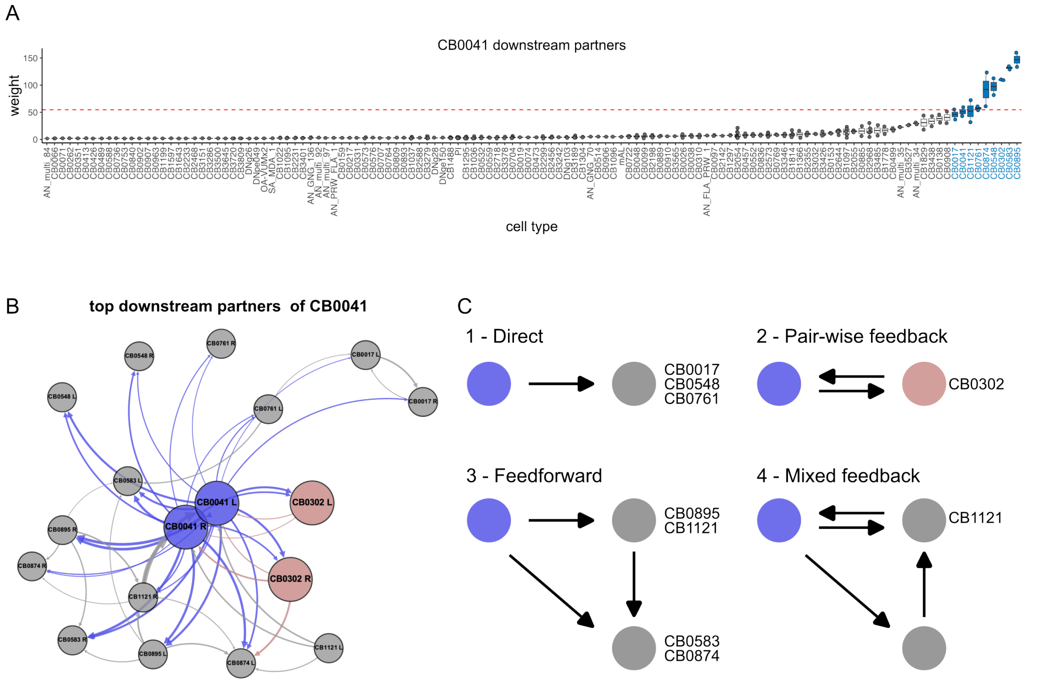


**Figure S13: CB0041 downstream partners and circuit motifs. A.** Connectivity between CB0041 and its downstream partners (absolute synaptic weights). Each filled circle represents a single downstream neuron. Boxes indicate the interquartile ranges. Black horizontal lines indicate the median value. Horizontal dashed lines (red) indicate the 95th percentile for the distribution of synaptic weights for all downstream partners. Cell types labeled with blue color are the top downstream partners, which have at least one neuron with a synaptic weight above the 95th percentile. **B.** Network representation of CB0041 and its top downstream partners. CB0302 is highlighted in salmon. **C.** Different network motifs between CB0041 and its downstream partners. CB0041 and CB0302 are connected with a pair-wise feedback motif.

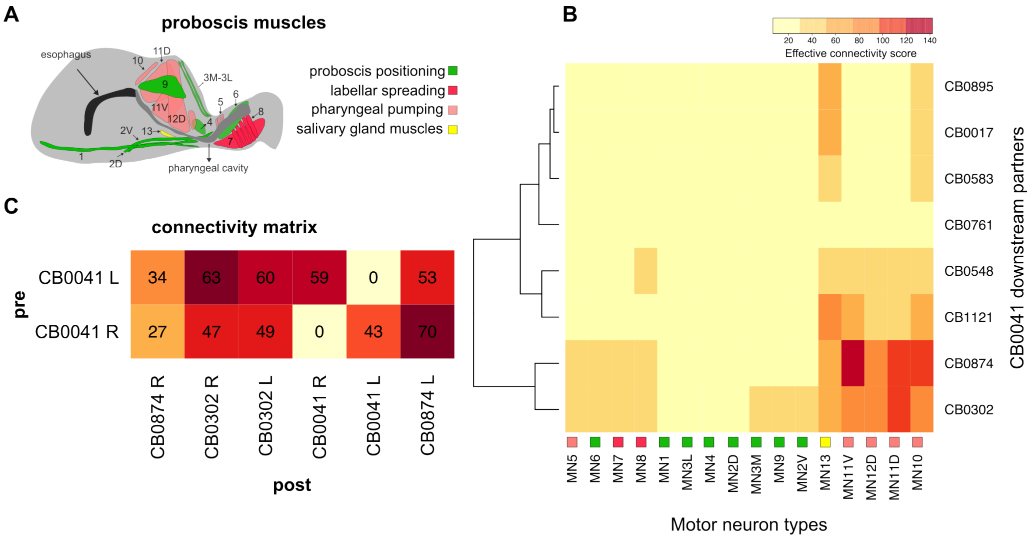


**Figure S14: CB0041 downstream partners and their effective connectivity with different motor neurons. A.** Proboscis muscle groups responsible for the movement of the proboscis during proboscis extension-retraction (labellum, haustellum, and rostrum) and ingestion (labellar spread, pharyngeal pumping, and salivation). Muscle groups are color-coded according to their function. **B.** Hierarchical clustering of top CB0041 downstream partners using effective connectivity scores with different motor neuron types. Motor neurons are color-coded according to the function they are involved in. CB0302 and CB0874 have relatively high effective connectivity scores with motor neurons involved in ingestion. **C.** Absolute synaptic weights between CB0041 and its downstream partners CB0302 and CB0874.

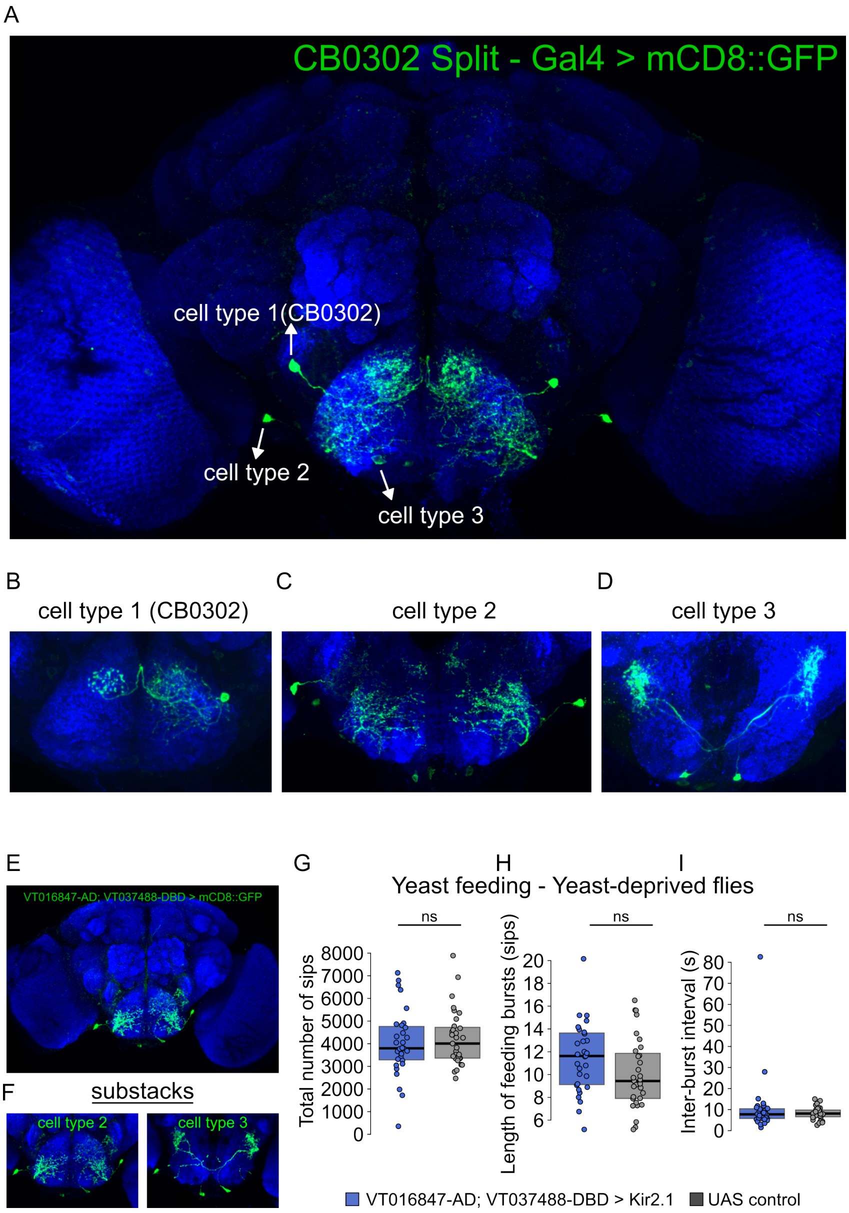


**Figure S15: Anatomical characterization of other cell types labelled by the Sustain Split-Gal4 driver line and excluding their role in controlling the feeding microstructure. A.** The expression pattern of the Split-Gal4 driver line used to label CB0302 (maximum intensity projection). Three cell types are labeled by the driver line as indicated in **B-D.** green: neurons expressing mCD8::GFP, blue: neuropile staining (nc82). Cell type 1 is CB0302. **E.** and **F.** Expression pattern of another Split-Gal4 driver line labeling both cell types 2 and 3. **E.** Maximum intensity projection of a full stack and **F.** substacks of E. **G-I.** Silencing cell types 2 and 3 simultaneously does not affect any feeding parameters. UAS control: *Empty Split-Gal4* x *UAS-Kir2.1*. Filled circles indicate individual flies. Boxes indicate the interquartile ranges. Horizontal lines indicate the median value. ns: not statistically significant, Wilcoxon rank-sum test.

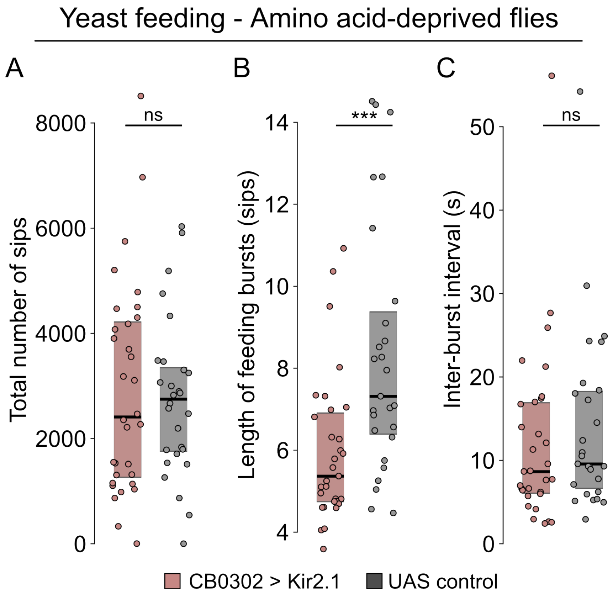


**Figure S16: Effect of silencing Sustain on feeding microstructure parameters in amino acid-deprived flies. A-C.** Silencing CB0302 in amino acid-deprived flies leads to **B.** shorter yeast feeding bursts, while not affecting **A.** the total number of yeasts sips or **C.** inter-sip intervals. UAS control: *Empty Split-Gal4* x *UAS-Kir2.1*. Filled circles indicate individual flies. Boxes indicate the interquartile ranges. Horizontal lines indicate the median value. ns: not statistically significant, Wilcoxon rank-sum test. ***: $p < 0.001$.

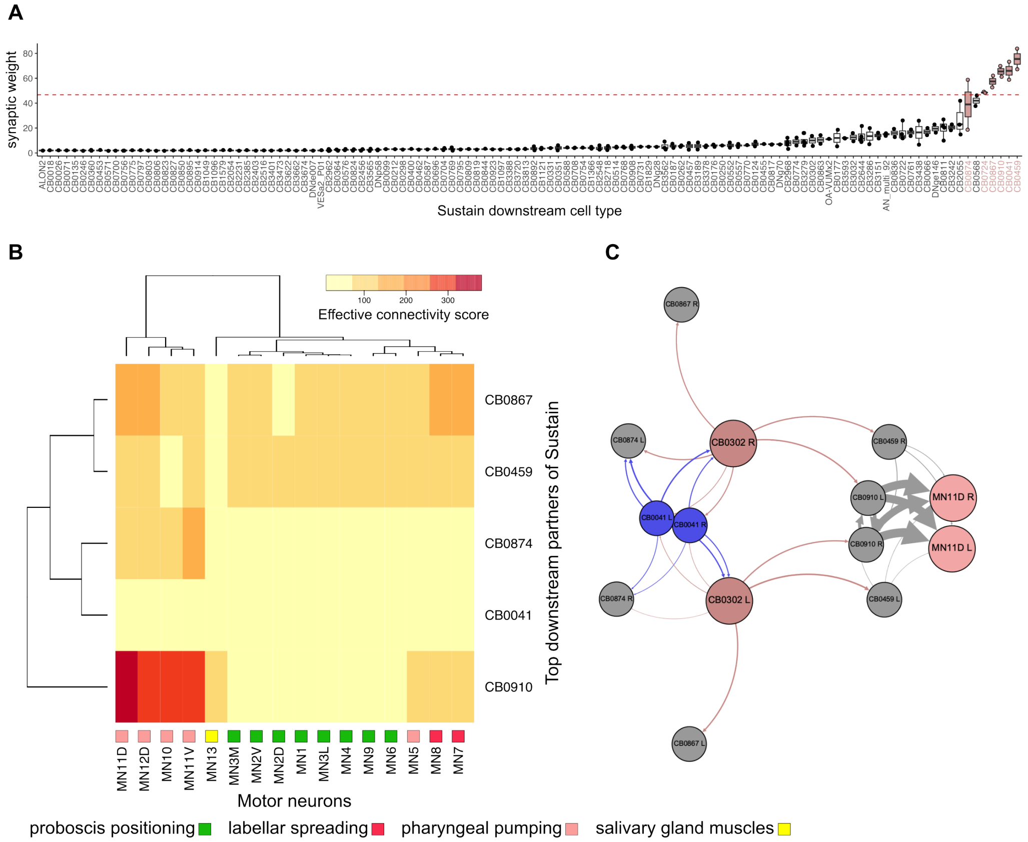


**Figure S17: Sustain downstream partners and their connectivity with proboscis motor neurons. A.** Connectivity between Sustain (CB0302) and its downstream partners (absolute synaptic weights). Each filled circle represents a single downstream neuron. Boxes indicate the interquartile ranges. Black horizontal lines indicate the median value. Horizontal dashed lines (red) indicate the 95th percentile for the distribution of synaptic weights for all downstream partners. Cell types labeled with light pink color are the top downstream partners, which have at least one neuron with a synaptic weight above the 95th percentile. **B.** Hierarchical clustering of top Sustain downstream partners using effective connectivity scores with different motor neuron types. Motor neurons are color-coded according to the function they are involved in. CB0910 has relatively high effective connectivity scores with pharyngeal motor neurons. **C.** Network representation of the circuitry connecting Sustain to MN11D.

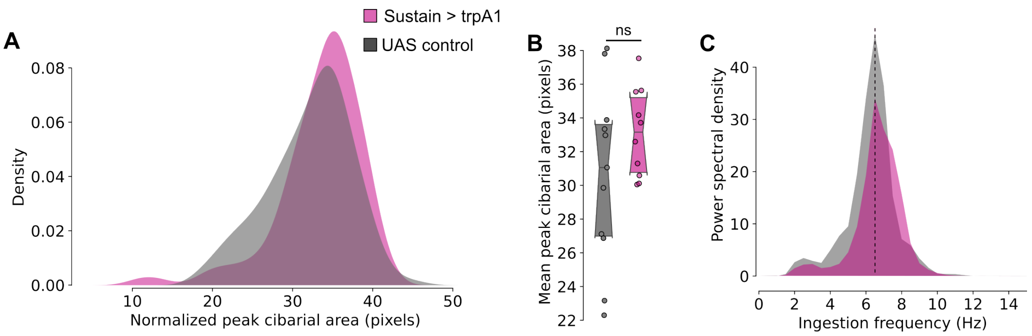


**Figure S18: Thermogenetic activation and its effect on pharyngeal pumping. A.** and **B.** Acute thermogenetic activation of Sustain using TrpA1 leads to a clear but statistically non-significant increase in ingestion volume. **A.** Kernel density estimations of the normalized peak cibarial area. **B.** Mean normalized peak cibarial area for individual flies. Boxes indicate the interquartile ranges. Horizontal lines indicate the median value. Notches indicate 95% confidence intervals. **C.** Power-spectral density of swallowing for Sustain-activated and control flies. Dashed vertical lines indicate peak ingestion frequency. UAS control: *Empty Split-Gal4* x *UAS-trpA1*. ns: not statistically significant.

| Genotype | Source | Figure | Supplementary Figure | Note |
|---|---|---|---|---|
| w[*]; P{w[+mC]=Ir76b-GAL4.916}226.8; TM2/TM6B, Tb[+] | BDSC#41730 | 1d,e | - | - |
| w[1118]; P{y[+t7.7] w[+mC]=GMR57F03-GAL4}attP2 | BDSC#46386 | 1f,g | - | - |
| w; 81E10-p65ADZp in attP40; VT023745-ZpGDBD in attP2 | Janelia Flylight | 4d,e,f,g | - | SS46917 (Rattle) |
| w; R81E10-p65ADZp in attP40; VT037804-ZpGDBD in attP2 | Janelia Flylight | - | 10e-j | SS31333 (Fdg) |
| w; 25A01-p65ADZp in attP40; 37D11-ZpGDBD in attP2 | Janelia Flylight | - | 10e-j | SS31386 (Bract) |
| w; 23G11-p65ADZp in attP40; VT003236-ZpGDBD in attP2 | Janelia Flylight | - | 10k-p | SS47744 (Roundup) |
| w; 11B11-p65ADZp in attP40; VT003236-ZpGDBD in attP2 | Janelia Flylight | - | 10k-p | SS47745 (Roundup) |
| w; 13F04-p65ADZp in attP40; 20E07-ZpGDBD in attP2 | This study | 4h,i,j,k | 12b,c | Split combination of BDSC#68831 and BDSC#69640 |
| w; VT010670-p65ADZp in attP40; VT037488-ZpGDBD in attP2 | This study | 5d,e,f,g; 7e,f,g | 15a-d; ; 16a-c; 18 | Split combination of BDSC#71246 and BDSC#74858 |
| w; VT016847-p65ADZp in attP40; VT037488-ZpGDBD in attP2 | This study | - | 15e-i | Split combination of BDSC#74216 and BDSC#74858 |
| w; VT020737-p65ADZp in attP40; 10B11-ZpGDBD in attP2 | Julie Simpson | 6d,e,f | - | MN11D, https://doi.org/10.7554/eLife.54978 |
| w[1118]; P{y[+t7.7] w[+mC]=p65.AD.Uw}attP40; P{y[+t7.7] w[+mC]=GAL4.DBD.Uw}attP2 | BDSC#79603 | 4e,f,g,i,j,k; 5d-g; 6e,f; 7e,f | 10a-f; h-t; 11; 12b,c; 15g-i; 16a-c | Empty Split-Gal4 |
| w;+; 5xUAS-Kir2.1:EGFP | Ribeiro Lab | 1e,f,g; 4e-g, i-k; 5d-g; 6e,f; 7e-g | 10a-f; h-t; 11; 12b,c | Recombined with w;+; tubGal80ts |
| w;+; tubGal80ts | Ribeiro Lab | 1e,f,g | - | Recombined with w;+; 5xUAS-Kir2.1:EGFP |
| CantonS | Ribeiro Lab | - | 1 | - |
| w; VT061715-p65ADZp in attP40; VT005008-ZpGDBD in attP2 | Julie Simpson | - | 11 | MN9, https://doi.org/10.7554/eLife.54978 |
| w; UAS-trpA1 | This study | - | 18 | - |

**Supplemental Table 1 : Fly genotypes used in this study.**

| Connectome | Body_ID | Root_Side | Type | Entry_Nerve | FAFB_version |
|---|---|---|---|---|---|
| FAFB – Flywire | 720575940622358057 | R | dorsal_tpGRN | MxLbN | 783 |
| FAFB – Flywire | 720575940625752778 | R | dorsal_tpGRN | MxLbN | 783 |
| FAFB – Flywire | 720575940627397263 | R | dorsal_tpGRN | MxLbN | 783 |
| FAFB – Flywire | 720575940630229007 | R | dorsal_tpGRN | MxLbN | 783 |
| FAFB – Flywire | 720575940631106664 | R | dorsal_tpGRN | MxLbN | 783 |
| FAFB – Flywire | 720575940632341452 | R | dorsal_tpGRN | MxLbN | 783 |
| FAFB – Flywire | 720575940611018714 | L | dorsal_tpGRN | MxLbN | 783 |
| FAFB – Flywire | 720575940622236671 | L | dorsal_tpGRN | MxLbN | 783 |
| FAFB – Flywire | 720575940625615705 | L | dorsal_tpGRN | MxLbN | 783 |
| FAFB – Flywire | 720575940616971035 | L | dorsal_tpGRN | MxLbN | 783 |
| FAFB – Flywire | 720575940606254002 | L | dorsal_tpGRN | MxLbN | 783 |
| FAFB – Flywire | 720575940604852146 | L | claw_tpGRN | MxLbN | 783 |
| FAFB – Flywire | 720575940607500297 | L | claw_tpGRN | MxLbN | 783 |
| FAFB – Flywire | 720575940608296917 | L | claw_tpGRN | MxLbN | 783 |
| FAFB – Flywire | 720575940609533230 | L | claw_tpGRN | MxLbN | 783 |
| FAFB – Flywire | 720575940610474610 | L | claw_tpGRN | MxLbN | 783 |
| FAFB – Flywire | 720575940611184819 | L | claw_tpGRN | MxLbN | 783 |
| FAFB – Flywire | 720575940613080751 | L | claw_tpGRN | MxLbN | 783 |
| FAFB – Flywire | 720575940618466384 | L | claw_tpGRN | MxLbN | 783 |
| FAFB – Flywire | 720575940620332404 | L | claw_tpGRN | MxLbN | 783 |
| FAFB – Flywire | 720575940620698136 | L | claw_tpGRN | MxLbN | 783 |
| FAFB – Flywire | 720575940622313971 | L | claw_tpGRN | MxLbN | 783 |
| FAFB – Flywire | 720575940622604109 | L | claw_tpGRN | MxLbN | 783 |
| FAFB – Flywire | 720575940623173470 | L | claw_tpGRN | MxLbN | 783 |
| FAFB – Flywire | 720575940624311881 | L | claw_tpGRN | MxLbN | 783 |
| FAFB – Flywire | 720575940624352643 | L | claw_tpGRN | MxLbN | 783 |
| FAFB – Flywire | 720575940624792197 | L | claw_tpGRN | MxLbN | 783 |
| FAFB – Flywire | 720575940625507870 | L | claw_tpGRN | MxLbN | 783 |
| FAFB – Flywire | 720575940625741114 | L | claw_tpGRN | MxLbN | 783 |
| FAFB – Flywire | 720575940626284036 | L | claw_tpGRN | MxLbN | 783 |
| FAFB – Flywire | 720575940627036746 | L | claw_tpGRN | MxLbN | 783 |
| FAFB – Flywire | 720575940627110672 | L | claw_tpGRN | MxLbN | 783 |
| FAFB – Flywire | 720575940629270339 | L | claw_tpGRN | MxLbN | 783 |
| FAFB – Flywire | 720575940631020079 | L | claw_tpGRN | MxLbN | 783 |
| FAFB – Flywire | 720575940631423263 | L | claw_tpGRN | MxLbN | 783 |
| FAFB – Flywire | 720575940633634199 | L | claw_tpGRN | MxLbN | 783 |

| Connectome | Body_ID | Root_Side | Type | Entry_Nerve | FAFB_version |
|---|---|---|---|---|---|
| FAFB – Flywire | 720575940633733601 | L | claw_tpGRN | MxLbN | 783 |
| FAFB – Flywire | 720575940635176366 | L | claw_tpGRN | MxLbN | 783 |
| FAFB – Flywire | 720575940635425893 | L | claw_tpGRN | MxLbN | 783 |
| FAFB – Flywire | 720575940635791167 | L | claw_tpGRN | MxLbN | 783 |
| FAFB – Flywire | 720575940635990896 | L | claw_tpGRN | MxLbN | 783 |
| FAFB – Flywire | 720575940638682787 | L | claw_tpGRN | MxLbN | 783 |
| FAFB – Flywire | 720575940608295689 | R | claw_tpGRN | MxLbN | 783 |
| FAFB – Flywire | 720575940608356053 | R | claw_tpGRN | MxLbN | 783 |
| FAFB – Flywire | 720575940609306563 | R | claw_tpGRN | MxLbN | 783 |
| FAFB – Flywire | 720575940610649442 | R | claw_tpGRN | MxLbN | 783 |
| FAFB – Flywire | 720575940611106453 | R | claw_tpGRN | MxLbN | 783 |
| FAFB – Flywire | 720575940611678181 | R | claw_tpGRN | MxLbN | 783 |
| FAFB – Flywire | 720575940612582514 | R | claw_tpGRN | MxLbN | 783 |
| FAFB – Flywire | 720575940616163701 | R | claw_tpGRN | MxLbN | 783 |
| FAFB – Flywire | 720575940616264788 | R | claw_tpGRN | MxLbN | 783 |
| FAFB – Flywire | 720575940619103435 | R | claw_tpGRN | MxLbN | 783 |
| FAFB – Flywire | 720575940619155403 | R | claw_tpGRN | MxLbN | 783 |
| FAFB – Flywire | 720575940619793940 | R | claw_tpGRN | MxLbN | 783 |
| FAFB – Flywire | 720575940622585224 | R | claw_tpGRN | MxLbN | 783 |
| FAFB – Flywire | 720575940622743309 | R | claw_tpGRN | MxLbN | 783 |
| FAFB – Flywire | 720575940623843145 | R | claw_tpGRN | MxLbN | 783 |
| FAFB – Flywire | 720575940624310676 | R | claw_tpGRN | MxLbN | 783 |
| FAFB – Flywire | 720575940626703241 | R | claw_tpGRN | MxLbN | 783 |
| FAFB – Flywire | 720575940628616271 | R | claw_tpGRN | MxLbN | 783 |
| FAFB – Flywire | 720575940630702789 | R | claw_tpGRN | MxLbN | 783 |
| FAFB – Flywire | 720575940632302161 | R | claw_tpGRN | MxLbN | 783 |
| FAFB – Flywire | 720575940632467923 | R | claw_tpGRN | MxLbN | 783 |
| FAFB – Flywire | 720575940633455073 | R | claw_tpGRN | MxLbN | 783 |
| FAFB – Flywire | 720575940637650522 | R | claw_tpGRN | MxLbN | 783 |
| FAFB – Flywire | 720575940639555928 | R | claw_tpGRN | MxLbN | 783 |
| FAFB – Flywire | 720575940644157988 | R | claw_tpGRN | MxLbN | 783 |
| FAFB – Flywire | 720575940645611652 | R | claw_tpGRN | MxLbN | 783 |
| FAFB – Flywire | 720575940646736772 | R | claw_tpGRN | MxLbN | 783 |
| FAFB – Flywire | 720575940647084164 | R | claw_tpGRN | MxLbN | 783 |
| FAFB – Flywire | 720575940614208914 | R | claw_tpGRN | MxLbN | 783 |

**Supplemental Table 2 : FlyWire IDs for tpGRNs**

**Supplemental Video 1 : Feeding behavior of the fly.** Slowed down video ($\sim$ 5X) of a fly feeding on a patch of yeast in a flyPAD arena. Proboscis extension, labellar spreading, and proboscis retraction are observed.

**Supplemental Video 2 : The pharyngeal pumping assay.** Pharyngeal pumping is quantified using DeepLabCut by tracking multiple points on the labellum, bristles, cibarium, and feeding capillary. A dyed food is used to visualize cibarial filling and emptying. Proboscis extension, labellar spreading, and pharyngeal pumping are observed.
